# New perspectives on multilocus ancestry informativeness

**DOI:** 10.1101/273466

**Authors:** Omri Tal, Tat Dat Tran

## Abstract

We present an axiomatic approach for *multilocus informativeness* measures for determining the amount of information that a set of polymorphic genetic markers provides about individual ancestry. We then reveal several surprising properties of a decision-theoretic based measure that is consistent with the set of proposed criteria for *multilocus informativeness*. In particular, these properties highlight the interplay between information originating from population priors and the information extractable from the population genetic variants. This analysis then reveals a certain deficiency of *mutual information* based *multilocus informativeness* measures when such population priors are incorporated. Finally, we analyse and quantify the inevitable inherent decrease in *informativeness* due to learning from finite population samples.

## 1 Introduction

> *“The investigations on the foundations of geometry suggest the problem: To treat in the same manner, by means of axioms, those physical sciences in which mathematics plays an important part.”*
>
> — – Hilbert 1902, p.454

The literature on inferring individual ancestry from genetic markers and the related measures of *ancestry informativeness* is vast and involves multiple perspectives and mathematical approaches (e.g. [Rosenberg et al., 2003]; [Ding et al., 2011]). The inferential task is conceptually linked to the *uncertainty* inherent in any effective classification scheme given a target set of polymorphic genetic markers from a number of source populations. However, *informativeness* is commonly interpreted a *property of the data* independent of any particular classification heuristic; in that sense it is ideally meant to capture only the *relevant* aspect of informational content – and this is why the formulation of a good measure is hardly straightforward. In this paper we adopt an essentially axiomatic approach which relies on first producing a set of appropriate and justifiable criteria that any measure should comply with. This is in contrast to previous approaches that have considered measures of *informativeness* in a more ad-hoc fashion, without anchoring them in any rigorous framework. Moreover, following the successful approach of [Shannon, 1948] in formalizing the transfer of information it is commonly recognized that the justification for regarding a quantity an information measure resides in the associated mathematical theorems demonstrating operational significance ([Csiszár, 2008]). Here we aim to abide to this realization by deriving novel properties of *ancestry informativeness* that are also of potentially practical significance.

This paper follows in the footsteps of the preliminary analysis laid out in [Tal, 2012b]. That work focused on reviewing and comparing between multiple candidates for *informativeness* based on simple distribution divergences (e.g. the class of *f-divergences*), distance metrics (the *Mahalanobis distance*) and differentiation measures (the population-genetic *F_ST_*) measures, and illuminating the drawbacks of each in in the context of a particular set of criteria. Here, we both refine and extend that set, and focus on information-theoretic and decision-theoretic *informativeness* measures, which are frequently invoked in the literature and also utilized in practical inference applications. Existing approaches for deriving measures for inferring ancestry have mostly focused on single-locus measures, have not rigorously incorporated the diminishing effects on *informativeness* of noise resulting from finite samples, and have not appropriately accounted for the effects of source-population size discrepancies. Crucially, previous related work has not sought justification in a firm conceptual or mathematically formal framework.

Our model highlights the important aspects of *ancestry informativeness* given simplifying assumptions on the nature of the genetic data. Theoretical work in population genetics often utilizes *haploid* rather than *diploid* models for the sake of simplified analysis (e.g. [Carja and Feldman, 2012]). We first consider a model of *haploid* populations given known allele frequencies from *biallelic* loci from two subpopulations with known class priors. The priors represent the discrepancy in source populations size, an aspect often incorporated in population models ([Rosenberg et al., 2003]; [Rosenberg, 2005]). Subsequently, sampling considerations enter into the analysis, and we derive enhanced *informativeness* measures that reflect more practical research studies.

## 2 The Population Model

We consider for simplicity a basic model of two haploid populations, denoted *P* and *Q*, a set of biallelic variants from these populations (following [Tal, 2012b]). We denote by *C_n_* an *informativeness* measure for ancestry inference across a set of *n* loci, which captures the information given by a set of polymorphisms and the knowledge of relative population sizes, and which crucially *complies with a given stipulated set of criteria*.

More rigorously, *C_n_*(*α, P, Q*) is a measure of *ancestry informativeness* given a set of *n* biallelic markers (such as SNPs), where here *P* and *Q* are *vectors* of known allele frequencies (*p*_1_, … , *p_n_*) and (*q*_1_, … , *q_n_*) from the respective populations, with 0 < *p_i_* < 1 and 0 < *q_i_* < 1. We (naturally) assume that *p_i_* and *q_i_* are true population parameters of *polymorphic* loci, i.e., each locus in each population is properly biallelic (the degrading effect of utilizing frequency *estimates* is introduced at a later section). A *genotype* sample of length *n* from one of the populations is then defined by an ordered sequence of *n* polymorphic alleles from the respective population, with a population frequency prescribed by the corresponding allele frequencies. We conveniently signify complete *informativeness* by an upper bound of 1, such that *C_n_*(*α, P,Q*) → 1 whenever asymptotically definite classification is inherently possible.

We differentiate between a *trivial* locus and an *uninformative* one. The former case involves allele frequencies that exactly equal between the two populations *p_i_* = *q_i_*, while the latter case the differentiation at that locus is below a threshold such that there is no contribution to *C_n_*. We shall show that there exist non-trivial but uninformative loci, i.e. that inclusion of loci with frequency differences greater than zero does not always contribute to *informativeness*.

The model includes a population prior *α*, which is arbitrarily assigned to population *P*, such that 1 − *α* is the prior of population *Q*. This prior is interpreted as the probability that a sample belongs to population *P* when its genotype is unknown, and simply reflect the known discrepancy in population sizes, treated from a Bayesian perspective, as in the model of [Rosenberg et al., 2003]. In effect, if we denote by *N_X_* the size of population *X*, then *α* = *N*_1_/(*N*_1_ + *N*_2_). Although the full notation is *C_n_*(*α,P,Q*), we will interchangeably use *C_n_* for simplicity in notation, where contextually sufficient.

Formally, let *X* = {0,1} be a binary variable representing the source populations *P* and *Q* respectively where, *X* ~ *Bernoulli*( 1 − *α*). Now let *Y_i_* = {0,1} be an allele at biallelic haploid locus *i*, with *p_i_* = *Pr*(*Y_i_* = 1|*X* = 0) and *q_i_* = *Pr*(*Y_i_* = 1|*X* = 1) where,

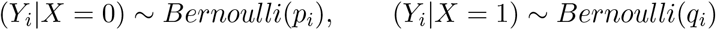

where the genotype frequencies assuming linkage equilibrium, *h_k_* for population *P* and *g_k_* for population *Q*, are a simple product of allele frequencies (formulated in a closed-form as in [Tal, 2012b]),

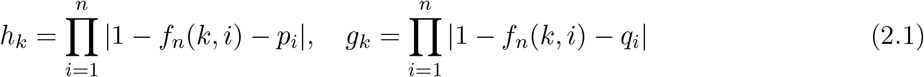

where

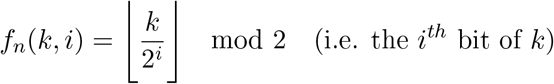

(e.g., for *n* = 3, *h*_0_ = (1 − *p*_1_)(1 − *P*_2_)(1 − *P*_3_), *h*_1_ = *p*_1_(1 − *p*_2_)(1 − *p*_3_), … ,*h*_7_ = *p*_1_*p*_2_*p*_3_).

### 2.1 The Criteria for *Informativeness* Measures

Here we specify a set of criteria for *multilocus informativeness* with justification stemming from established empirical studies and theoretical results, and from basic intuitive reasoning (subsequently elaborated on in the *discussion*). This set of criteria are an *extended, corrected* and *refined* reformulation following the preliminary treatment in [Tal, 2012b].

1. *Zero: C_n_* = 0 *if and only if* the two populations are virtually the same population, i.e., across *n* loci, *p_i_* = *q_i_* for all *i* and (implicitly) the prior 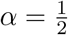. Naturally, under a single class (population) the classification task becomes void given any set of loci, while at the other hand, any level of population structure captured by a set of loci should admit some potential for successful assignment.
2. *Performance: C_n_* should be a monotonic non-decreasing function of *n*. Informally, each additional locus should *potentially* be informative given the set of loci already included in C_n_, but never diminish the aggregated informativeness. This criterion is justified by the phenomenon of *asymptotically perfect classification* which is achievable when effectively utilizing information from the intrinsicly high dimensional nature of functional data ([Delaigle and Hall, 2012]).
3. *Asymptotics: if* allele frequencies at each locus differ between populations by *at least ε* > 0, where *ε* is any *predefined* value as small as we wish, then *C_n_* → 1 as *n* → ∞. Informally, with this *C_n_* an infinite number of loci with even very slight frequency differences between the two populations induces complete informativeness. This criterion is strongly justified from both empirical studies ([Witherspoon et al., 2007]) with high-sequencing data and from theoretical considerations on separation in high dimensional spaces ([Edwards, 2003], [Tal, 2012a]).
4. *Neutrality*: The inclusion of *trivial loci* (*p_i_* = *q_i_*) should not affect *C_n_*, as naturally the two populations are not differentiated with respect to such loci. Note that the alleles at such loci are still polymorphisms within each population, but these polymorphisms occur with (theoretically) equal frequencies.
5. *Continuity: C_n_* should be continuous in *p_i_, q_i_* and *α*,, as one would not expect small fluctuations in allele frequencies at any locus or in population sizes to have large effects on informativeness.
6. *Dominance*: for any finite number of loci *n*, we expect *C_n_* to be maximal if and only if for some locus *i* the differentiation is maximal, i.e. *δ_i_* = |*q_i_* − *p_i_*| → 1. Informally, any single locus with maximal allele frequency difference is sufficient for *accurate assignment* of any genotype: one may simply classify according to the presence or absence of a given allele at that locus. The ‘if and only if’ assures that (for any finite n) no other scenario that does not include *δ_i_* → 1 would result in *C_n_* → 1. The asymptotic limit (→ 1) here follows from the continuity criterion.
7. *Delta*: Overlapping allele-frequency differences should induce a strict ranking among loci. More precisely, when some locus *i* has a *wider and completely overlapping allele frequency difference* compared to locus *k* (without loss of generality, *p_i_* < *p_k_* and *q_i_* > *q_k_*), then the inclusion of locus *i* in *C_n_* should result in higher or equal total *informativeness* vs. the inclusion of *k*. Naturally, the contribution to *informativeness* of a marker which is both rarer in one population and more common in the other population – in relation to some other marker – should always be greater.
8. *Invariances*: Naturally, we expect *C_n_* to admit to several natural invariances and symmetries: [a] invariant to different ordering of sequenced loci, i.e., the components of the allele frequency vectors *P* and *Q* may be specified in any order, as long as they remain in synchrony; [b] symmetric with respect to the two populations, i.e., *C_n_*(*P, Q*) = *C_n_*(*Q, P*); [c] invariant to the arbitrary choice of the alleles to which we assign the frequency parameters – the simultaneous substitution of *p_i_* with (1 − *p_i_*) in *P* and *q_i_* with (1 − *q_i_*) in *Q*.
9. *Prior: C_n_* → 1 if *α* → 0 or *α* → 1, since if the discrepancy of source population size is extremely large, the probability for correct assignment should be asymptotically 1, irrespective of the allele frequency values. In that mostly hypothetical case, one would simply assign any unknown genotype to the large population. The use of limits here results from the continuity criterion and the framework that specifies *C_n_* < 1 and 0 < *α* < 1.

In formal terms:

Let *P* = (*p*_1_, … , *p_n_*), *Q* = (*q*_1_, … , *q_n_*), allele frequencies 0 < *p_i_,q_i_* < 1, population *P* prior 0 < *α* < 1. *C_n_*(*α, P, Q*), abbreviated *C_n_*(*P, Q*) or just *C_n_*, should satisfy:

1. *C_n_* = 0 iff *P* = *Q* (∀*i p_i_* = *q_i_*, 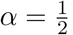)
2. *C*_*n*+1_ ≥ *C_n_*
3. ∀ *ε* > 0, if 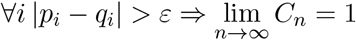
4. *p*_*n*+1_ = *q*_*n*+1_ ⇒ *C*_*n*+1_ = *C_n_*
5. *C_n_* is continuous in *p_i_, q_i_* and *α*
6. ∀*n, C_n_* → 1 iff |*p_i_* − *q_i_*| → 1 for some *i*
7. If without loss of generality, *p_n_* < *q_n_*, then for all *ε* : 0 < *ε* < 1 − *q_n_, C_n_*(*P*, (*q*_1_, … , *q_n_* + *ε*)) ≥ *C_n_*(*P, Q*) and for all *ε* : 0 < *ε* < *p_n_*, *C_n_*((*p*_1_, … , *p_n_* − *ε*), *Q*) ≥ *C_n_*(*P, Q*).
8. *C_n_*(*P, Q*) = *C_n_*((*p*_*σ*(1)_, … , *p*_*σ*(*n*)_, (*q*_*σ*(1)_, … , *q*_*σ*(*n*)_)) for all permutation *σ* ∈ *S_n_* *C_n_*(*α,P,Q*) = *C_n_*(1 − *α, Q,P*) *C_n_*(*P,Q*) = *C_n_*((*p*_1_, … , 1 − *p_i_*, … , *p_n_*), (*q*_1_, … , 1 − *q_i_, … , q_n_*))
9. 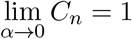 and 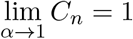

Obviously, any measure *C_n_* that complies with the set of criteria admits an infinite number of ‘correlated’ measures representing the degrees of freedom of *C_n_* (just as Shannon entropy *H* has degrees of freedom represented by a linear factor). Formally, this sense of a correlation between two functions *f* and *g* implies that for any two sets of parameters *x* and *y*,

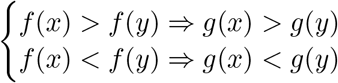

It is easy to show that equivalently, this implies the existence of a monotonic function *h* such that *f* = *h*(*g*). In terms of our inferential framework, this means that for any two panels of SNPs (along with corresponding population priors) represented by *x* and *y* above, correlated *informativeness* measures always admit the same ranking, and one is a monotonic function of the other. In Appendix F we prove that any measure correlated with *C_n_* also complies with our set of criteria.

### 2.2 Informativeness based on information-theoretic concepts

A well-known information theoretic measure of shared entropy is the *mutual information*, also commonly interpreted and utilized as a powerful measure of statistical dependency, sensitive also to nonlinear functional relationships ([Steuer et al., 2002]). The mutual information between an allele at a single locus and the source population has been explored in the context of *feature selection* ([Peng et al., 2005]) and *ancestry informativeness* ([Rosenberg et al., 2003]).

We would like to examine this instantiation of mutual information as a candidate for our Cn. From basic definitions of mutual information and conditional probability, we utilize our assumption of linkage equilibrium within each population to express the joint multivariate distributions [*Y*_1_, … , *Y_n_*|*X*] and [*Y*_1_, … ,*Y_n_*] in terms of the allele frequencies *p*(*y_i_*−*X*). With the population priors translating into *P*(*X* = 0) = *α, P*(*X* = 1) = 1 − *α*, we get (see [Tal, 2012b , Eq. 2]),

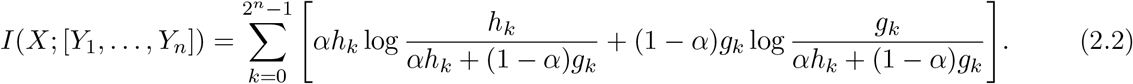

To comply with an upper bound of 1 corresponding to complete informativeness, normalization is required in the formulation of *C_n_*. The maximal value of this expression of mutual information is but since *H*(*X*) would be the minimum of the two in all non-trivial cases, we may normalize by *H*(*X*) = −*α* log *α* − (1 − *α*) log(1 − *α*),

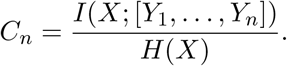

However, this candidate then fails the *zero* criterion (since *C_n_* = 0 if *p_i_* = *q_i_* for all values of prior α) and most crucially fails the *priors* criterion, since *C_n_* = 0 instead of 1 as *α* approaches 0 or 1, as illustrated in Fig. 1 (correcting Fig. 1B in [Tal, 2012b], which lacks proper normalization). The failure of the priors *criterion* is a characteristic of the non-normalized formulation as well, simply since *H*(*X*) is virtually zero at the prior extremes. Therefore, the deficiency exposed here similarly applies to the non-normalized *informativeness for assignment* measure, denoted *I_n_* from [Rosenberg et al., 2003]. We shall therefore henceforth refer to the mutual information-based informativeness as *I_n_*.

**Fig. 1:**
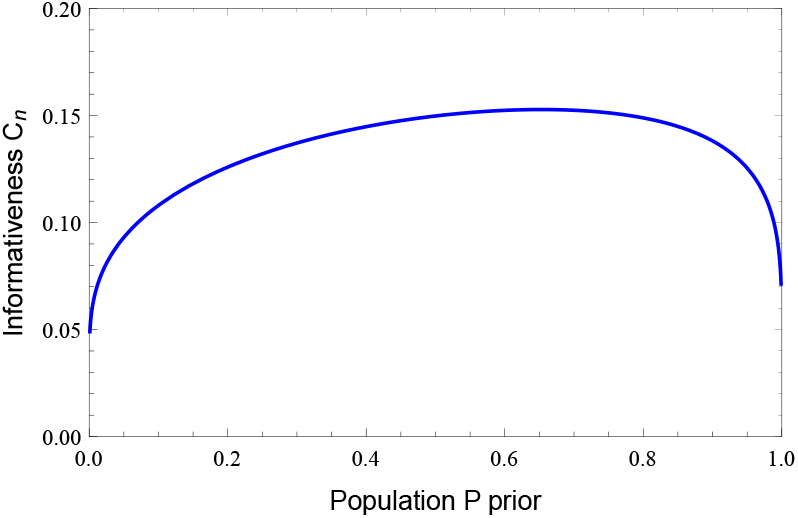
*C_n_* based on (normalized) *mutual information* fails the *priors* criterion. In this example, allele frequencies are *p*_1_ = 0.03/*q*_1_ = 0.18; *p*_2_ = 0.03/*q*_2_ = 0.18,*p*_3_ = 0.20/*q*_3_ = 0.48.

A related information theoretic measure is the *variation of information* (VI). It has been used as a criterion for comparing two partitions or clusterings of the same data set, and measures the amount of information lost and gained in changing from one cluster to another ([Meila, 2007]). The measure is defined as the difference between the joint entropy and the mutual information, and has the benefit of being a true metric.

Expressing VI between the population and genotype distributions in terms of our variables and incorporating the priors we have,

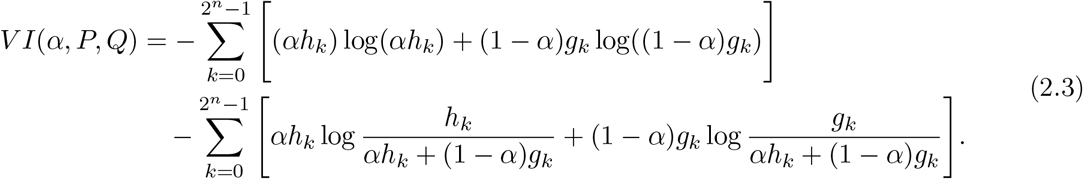

As with the previous candidate, we normalize by the maximal value attained by VI, which here is the joint entropy. Therefore,

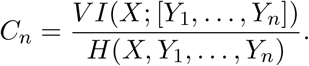

However, this candidate fails the important *dominance* criterion, irrespective of the normalization factor chosen (in fact, *C_n_ decreases* as the *absolute allele frequency difference* increases at a locus), as illustrated in Fig. 2.

**Fig. 2:**
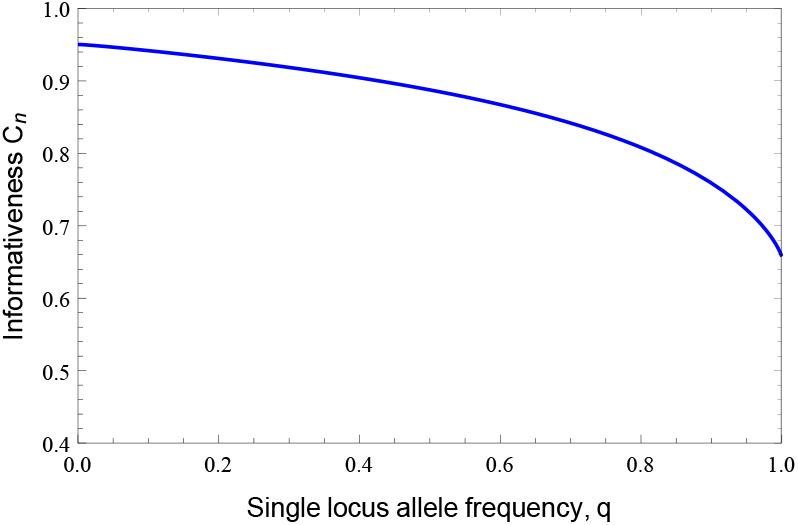
*C_n_* based on the *variation of information* fails the *dominance* criterion. In this example the population parameters are as in Fig. 1.

Some intuition about the deficiency of mutual information between the class and genotype in capturing *informativeness when population priors are incorporated* is gained by noting that mutual information represents the reduction of uncertainty in one random variable when knowing the other. Typically, in the ancestry inference framework that occupies us here, this means that *I*(*X; Y*) is the (average) *reduction of uncertainty about the source population X*, when *knowing* the genotype at *n* loci *Y* = [*Y*_1_, … , *Y_n_*]. By symmetry of mutual information, this inevitably also represents the *reduction of uncertainty about the* genotype at *n* loci when *knowing* the *source population X*, but since typically *H*(*Y*) ≫ *H*(*X*) for *n* ≫ 1, this reduction in *H*(*Y*) is relatively *inconsequential*. The absolute amount of reduction in source population uncertainty strictly depends on the initial amount of its uncertainty, which is encapsulated by the prior *ε*. This explains why the *priors* criterion fails with *I_n_*: as the prior approaches the extremes of 0 or 1, *H*(*X*) → 0 and consequently the reduction in this quantity also *approaches zero*. But this scenario represents *virtual certainty* about inferring the source population, such that *I_n_* should have approached 1 instead. This basic insight is schematically depicted in Fig. 3. The normalized formulation similarly fails this criterion, since,

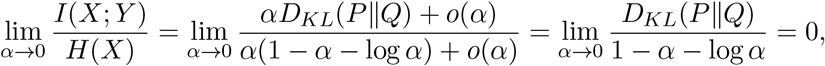

and similarly 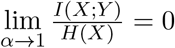

**Fig. 3:**
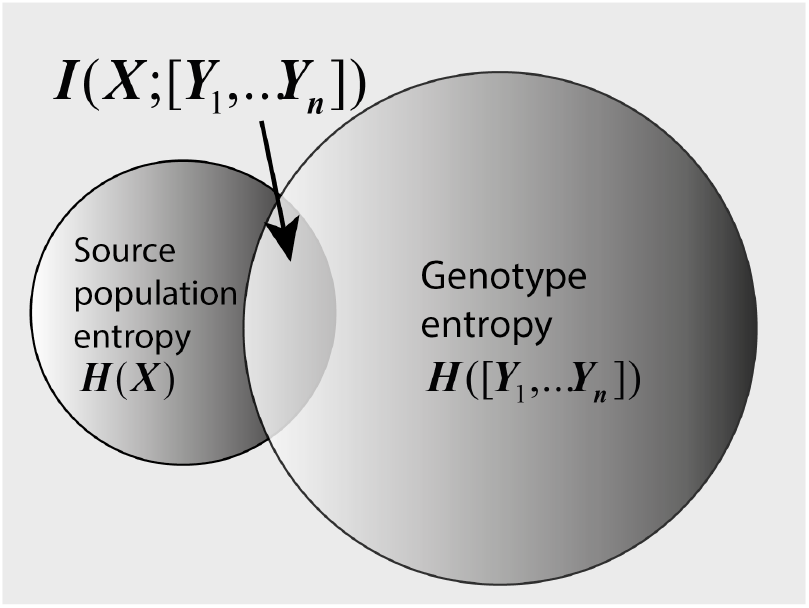
The reduction of uncertainty about the *source population X*, when *knowing* the genotype *Y* at *n* loci.

## 3 *C_n_* Based on a Naïve Bayes Classifier

The optimal classifier under known class-conditional densities is the Bayes *classifier (with class priors)* or alternatively the *maximum-likelihood* (ML) classifier, where data are classified to the most probable class. The Bayes error then is the expected error of this classifier, usually under a 0/1 risk function ([Hastie et al., 2009]). Here, we wish to derive a simple formulation of the Bayes error in the context of our population genetic model. The assumption of linkage equilibrium within each source population corresponds to within-class stochastic independence, and motivates the use of the popular a *naïve Bayes* classifier. The genotype frequencies are then simply the product of population-conditional allele frequencies across an independent set of loci (as in [Cornuet et al., 1999]; [Phillips et al., 2007]; [Tal, 2012b]). A similar approach for a decision-theoretic *informativeness* is the *multilocus* version of the *optimal rate of correct assignment* (ORCA) with general priors from [Rosenberg et al., 2003]. From basic definitions, the Bayes error for discrete data from two classes can be expressed as a prior-weighted sum of probabilities over all instances of the data ([Hastie et al., 2009]). For our framework, this translates to a sum over the 2^*n*^ possible genotypes, indexed by *k*,

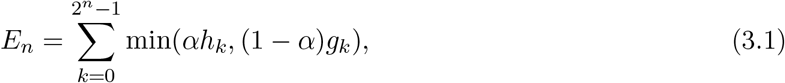

where the genotype frequencies *g_k_* and *h_k_* are defined in Eq.(2.1). Since for two classes the Bayes error is bounded below by 0 and above by 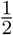, a straightforward transformation equivalent to proper normalization is,

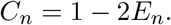

Thus,

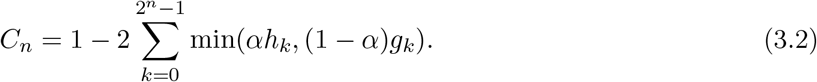

It is possible to derive an *equivalent* formulation based on the *variational distance*, a form of *f-divergence* ([Nguyen et al., 2009, section 2.1.1]), with a natural modification to incorporate our priors (see Appendix A for proof),

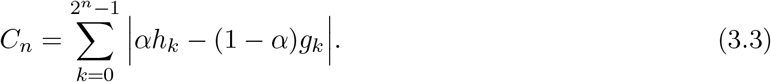

where the genotype frequencies are again as in Eq. (2.1).

Finally, we would like to produce a general formulation of *C_n_* which separates the *classifier function* (denoted here, with corresponding *indicator function*) from the error rate formalism. This will serve us in subsequent sections for deriving other decision-theoretic forms of *C_n_*. Following Eq. (3.2) we can write,

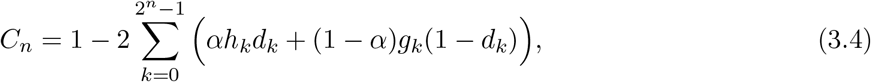

where

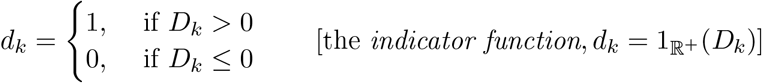

(in case of equal distances, *D_k_* = 0, we arbitrarily choose to classify to population *P*). For the *naïve Bayes*, *D_k_* simply compares the genotype probabilities weighed by the priors,

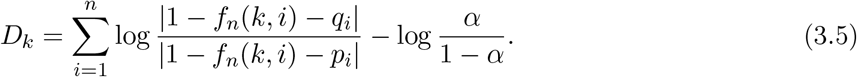

It is proven in Appendix B that these equivalent formulations of *C_n_* satisfy the complete set of *informativeness* criteria.

In fact, it is possible to derive a parametrized family of measures of infinite cardinality, based on a specific generalization of the *variational distance* that is also compliant with the full set of criteria (proof in Appendix G). Formally, denote this family of measures by 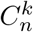, parametrized by the integer *k* : 1, … , ∞,

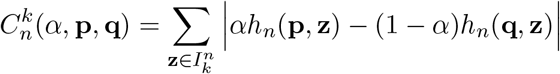

where **p** = (*p*_1_, … , *p_n_*), **q** = (*q*_1_, … ,*q_n_*), 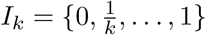 and

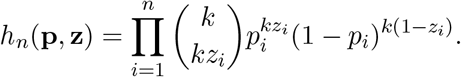

Note that 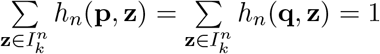.

Crucially, it can be shown that for any *k*, 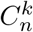 *is qualitatively different or uncorrelated with the Bayes-based C_n_*, i.e., is not a monotonic function of it (Appendix H). The compliance of the family of measures 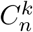 with all criteria therefore implies that the Bayes-error *C_n_* is not a *unique solution* in the context of our axiomatic framework for an *informativeness* measure.

The benefit of the Bayes-error *C_n_* over any instantiation from the infinite class 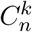 derives from the former’s *simplicity* in both formulation and computational cost in comparison to the latter. Therefore, we will henceforth focus our analysis on the simpler formulation.

We now reveal a core discrepancy of our Bayes-error *C_n_* and the information-theoretic *I_n_*. We first demonstrate that *C_n_* and *I_n_* are not correlated (i.e., no function *h* exists such that *I*_1_ = *h*(*C*_1_)) even for the most rudimentary case of *n* = 1 and equal priors. To see this, denote *I*_1_ by *f*(*x, y*) and *C*_1_ by *g*(*x,y*), both a function only of the two allele frequencies *x* and *y* at a single locus (setting *α* = 0.5), such that,

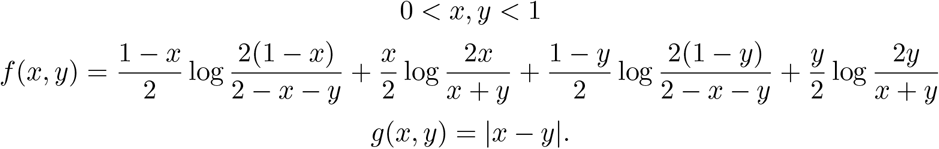

To prove *f* and *g* are not correlated we only need to identify two points (*p, q*) and (*p*′, *Q*′) such that *g*(*p, q*) = *g*(*p*′, *q*′) but *f*(*p, q*) ≠ *f*(*p*′, *q*′). It is easily verified that one such instance is (*p, q*) = (0.3,0.1) and (*p*′,*q*′) = (0.3, 0.5). This lack of correlation even for the most rudimentary case strongly implies non-correlation in the general sense of *n* ≥ 1. Crucially, simulations indeed indicate many cases of discrepancy between *C_n_* and *I_n_* in their ranking for *informativeness* of particular marker panels and associated values of the population prior. In such cases, the relative ranking given by *C_n_* or *I_n_* to these panels is almost surely prior-dependent (as illustrated by one such typical scenario in Fig. 4).

**Fig. 4:**
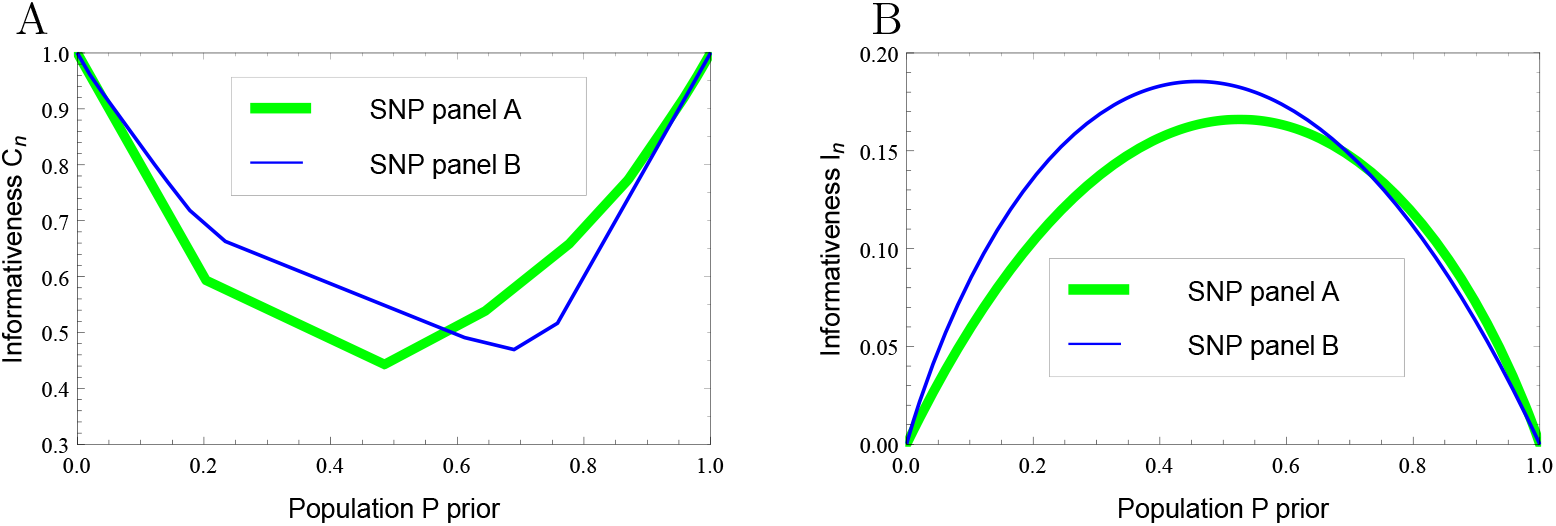
A simulation of one scenario demonstrating the discrepancy (curves intersect at different locations relative to the prior) between the decision-theoretic *C_n_* and the [Rosenberg et al., 2003] information-theoretic *I_n_*, as measures of *multilocus informativeness*. At a population prior of 0.65 *panel A* is deemed superior according to *C_n_* (Fig. 4A), while *panel B* is deemed superior according to *I_n_* (Fig. 4B). Both panels consist of 6 SNPs (*panel A*: 3 loci with allele frequencies *p* = 0.03/*q* = 0.18 and 2 loci with *p* = 0.20/*q* = 0.48; *panel B*: 3 loci with *p* = 0.30/*q* = 0.04 and 2 loci with *p* = 0.32/*q* = 0.25).

### 3.1 Several distinctive properties of the decision-theoretic informativeness

In this section we reveal insightful set of properties of the Bayes-based *C_n_*, which expecially highlight a unique balance between information originating from the genetic data vs. information attributable to population size discrepancy, represented by the priors (see Appendix C for the proofs of these properties).

a. *Loci-subadditivity (w/out priors)*: The sum of the informativeness of two sequences of loci cannot be lower than the informativeness of the aggregated sequence,

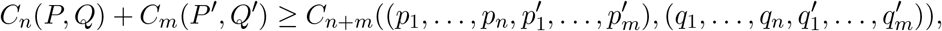

or put more simply in terms of an ordered lists concatenation operator ║,

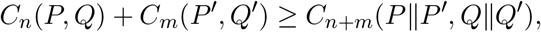 For *n* = *m* = 1 the strength of this subadditivity (the inequality above) reflects the redundancy in *informativeness* within a pair of individual loci. We note that when population priors are incorporated in *C_n_*, this property no longer necessarily holds.^1^ We may nevertheless allow for priors by introducing a correction term | 1 − 2*α*|,

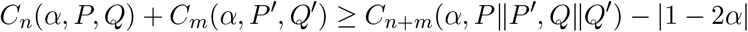

at the expense of overstretching somewhat the meaningful sense of proper subadditivity (see proof in Appendix C).
b. *Population-subadditivity (w/out priors): C_n_* complies with a triangle inequality between any three populations. Formally, for any *n* loci given three populations *P, Q and R*,

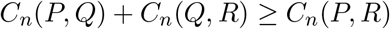 This property affords the interpretation of *informativeness* as a distance *metric* between populations, relative to any given number of loci (n). We note here that (contrary to the erroneous claim in [Tal, 2012b]) subadditivity does not generally hold with general priors,^2^

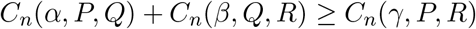

where the three priors are derived from relative population sizes *N*_1_,*N*_2_ and *N*_3_ of *P, Q* and *R* respectively such that, *α* = *N*_1_/(*N*_1_ + *N*_2_),β = *N*_2_/(*N*_2_ + *N*_3_),γ = *N*_1_/(*N*_1_ + *N*_3_) and thus any two priors define the third, e.g.,

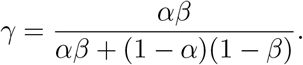
c. *Prior-washout*: The effect of the prior is *washed out* with additional loci; i.e., the effect on informativeness from the consideration of population priors *generally diminishes* with more loci. Formally, the range for *C_n_* as a function of *α* (0 < *α* < 1) never increases with *n* → *n* + 1, i.e. the minimal value of *C_n_*(*α*) increases or is unchanged (note this does not imply that for any two priors and any *n*, |*C*_*n*+1_(*α*_1_) − *C*_*n*+1_(*α*_2_)| ≤ |*C_n_*(*α*_1_) − *C_n_*(*α*_2_)|). Moreover, asymptotically with *n* the prior is completely washed out becoming uninformative.
d. *Prior-sensitivity: C_n_* as a function of the prior *α* is *convex* and the minimum is *unique almost surely* (and *C_n_* is not necessarily minimal at 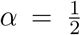). This means that *C_n_* is always sensitive to fluctuations in the prior.
e. *Prior-dominance*: There is a particular asymmetric balance between information stemming from the population priors and information from the genetic markers. For any given allele frequencies at *n* loci, there exist thresholds for prior disparity, beyond which the information provided by the priors eclipses the information provided by genetic markers. At that level of disparity and beyond, *C_n_* [a] is only a function of the prior *α*, [b] is insensitive to small fluctuations of allele frequencies, and [c] is ultimately unaffected by the exclusion of any or all *n* loci. In consequence, for any panel of markers there always exists a level of prior disparity (as a function of the allele frequencies) beyond which an effective classifier may ignore the given genotype sample, simply assigning it to the population with the larger prior.
f. *Uninformative-loci*: There always exists some such that the inclusion of an extra locus with an allele frequency difference does not change *C_n_*.^3^ Formally, for any *P* and *Q* with *n* loci, and for any prior *α*, without loss of generality,

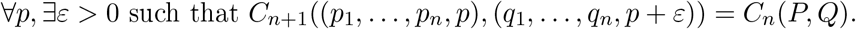 We would next like to inquire whether a formulation of *C_n_* based on the error rate of *any* valid classifier may also satisfy our axiomatic framework, or perhaps not all classifiers are equivalent in this sense.

## 4 *C_n_* based on a Nearest Centroid (NC) Classifier

Whereas a Bayes classifier is based on the probability of observing a given genotype in a target population, distance methods assign a genotype to the “closest” population ([Liao et al., 2009], [Degen et al., 2017]). In particular, we will require a genetic distance metric adopted as a measure between an individual and a population, or rather, the population genetic *centroid*. Importantly, these centroids are unique - defined irrespective of the *distance* measure one employs, and irrespective of the presence or absence of LD in the data or model.

In practical settings, the centroids are *learned* from the training data of m individual samples, *x*_1_, … , *x_m_* and computed using a simple arithmetic average: *M* = (*x*_1_ + ⋯ + *x_m_*)/*m*, (simulated in Fig. 5) or in terms of our model, *M* = [*E*(*Y*_1_|*X*), … , *E*(*Y_n_*|*X*)] such that, *M*_1_ = (*p*_1_, … , *p_n_*) and *M*_2_ = (*q*_1_, … , *q_n_*).

**Fig. 5:**
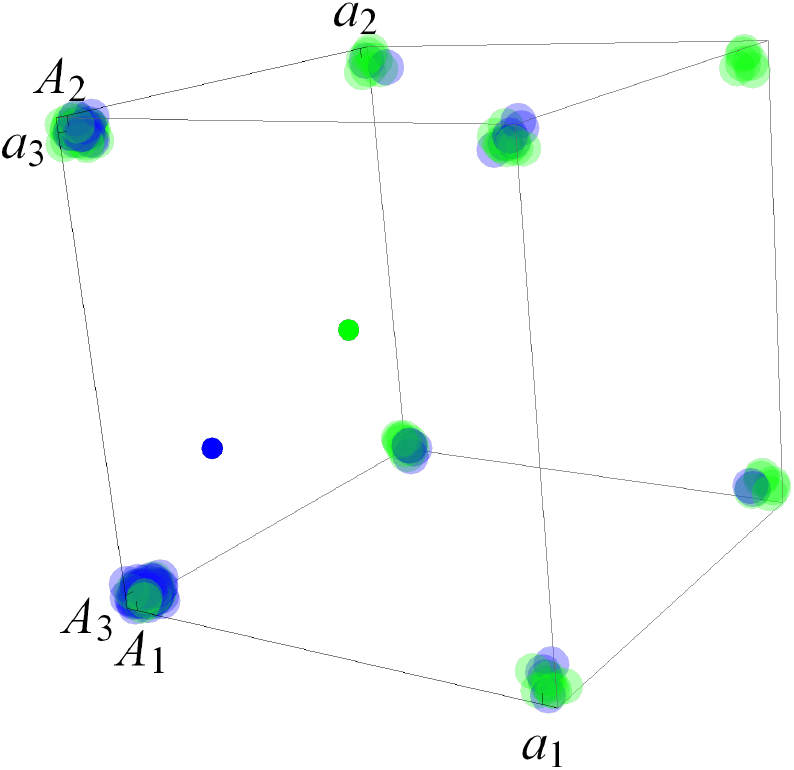
A 3–dimensional hypercube depiction of samples of three-locus genotypes from two haploid biallelic populations, and their centroids (blue and green). Each sphere at a vertex is an individual sample defined by alleles (a and A) at three loci; the small spheres represent population means. 80 samples from each population were randomly drawn from two populations with linkage equilibrium in each; allele frequencies are *p*_1_ = 0.2/*q*_1_ = 0.4; *p*_2_ = 0.1/*q*_2_ = 0.3; *p*_1_ = 0.3/*q*_2_ = 0.5.

The distance between any genotype *x_k_*, defined here as the *k*-th coordinate in {0,1}^*n*^, and centroid *M*_1_ depends on the distance metric we chose to employ. Using a *Euclidean* metric,

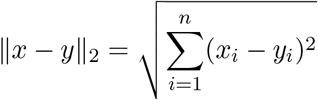

the distance to the centroid is,

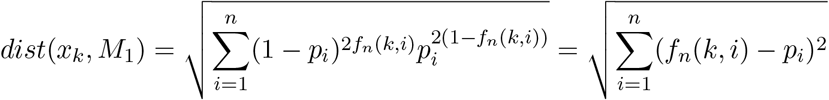

where as before, *f_n_*(*k,i*) := the *i^th^* bit of *k*. Alternatively, using a *Manhattan* distance or *L*_1_ norm,

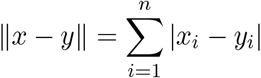

and the distance to the centroid would be,

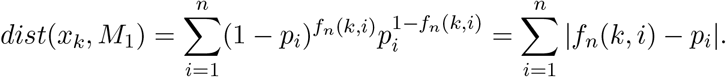

In the context of a genotypic space, the *L*_1_ norm is equivalent to the *allele-sharing-distance* (ASD). and is therefore more appropriate for our framework than the Euclidean, and we therefore confine our development to the former (the Euclidean distance is the square root of the ASD only for distances *between genotypes* but not for distances to *centroids*). The classifier term is then simply a difference of the distances of a genotype to the two centroids, i.e.,

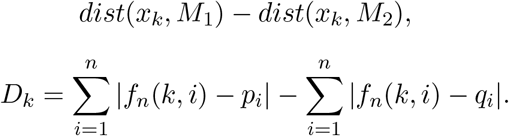

However, *C_n_* based on such classifier does not comply with the crucial *dominance* criterion for any *n* ≥ 3, as illustrated by counter-example in Fig. 6.

**Fig. 6:**
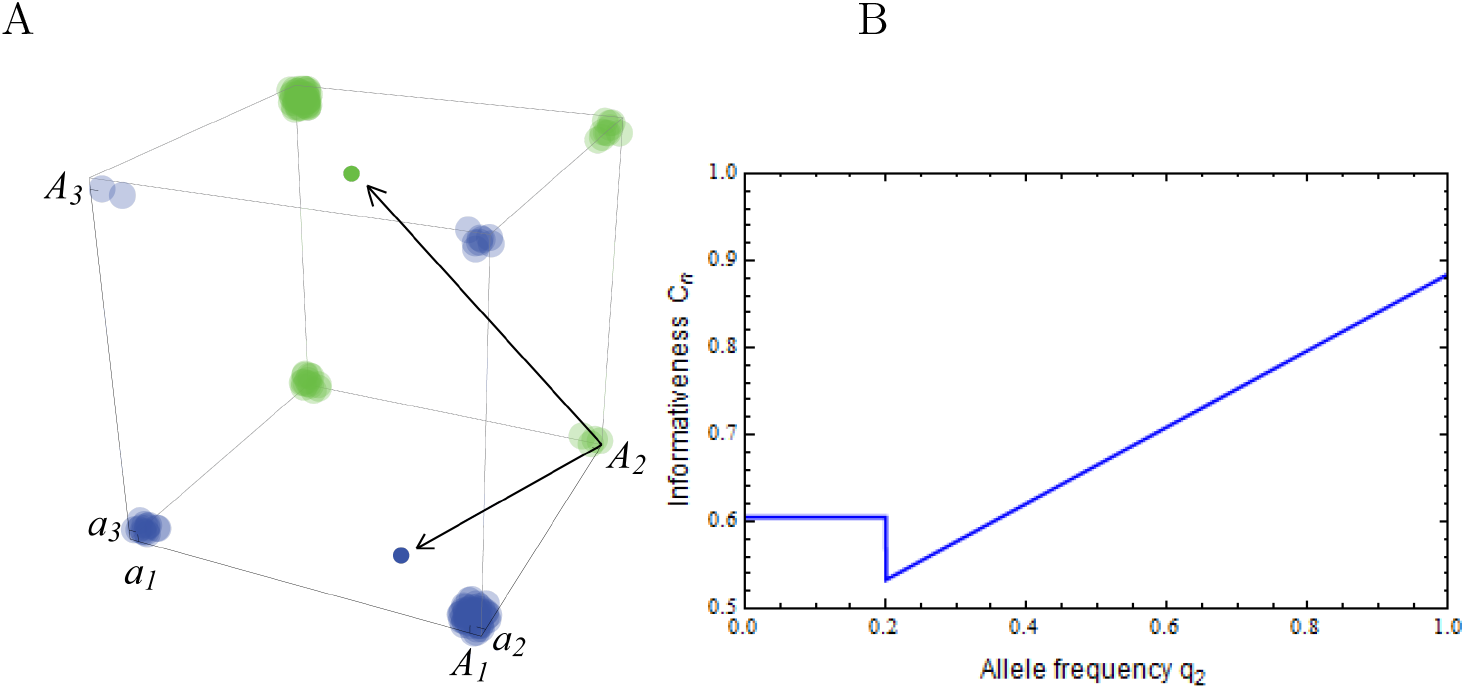
The *dominance* criterion fails for *C_n_* based on a nearest-centroid classifier (allele frequencies *p*_1_ = 0.8/*q*_1_ = 0.4, *p*_2_ ≈ 0/*q*_2_ ≈ 1,*p*_3_ = 0.2/*q*_3_ = 0.8). | A: A hypercube representation of a 3—loci scenario with 40 samples from each population; although there is complete separation along one dimension (*p*_2_ = 0, *q*_2_ = 1), some vertices may be wrongly classified (genotype *A*1*A*2*a*3 is closer in both *L*_1_ and *L*_2_ to the mean of the blue population, despite belonging to the green population). | B: *C_n_* < 1 despite reaching complete frequency divergence at locus #2 (*p*_2_ ≈ 0, *q*_2_ → 1).

The underlying problem is that the *L*_1_ metric does not take into account the variances of the genotype distribution of each population along the direction from any genotype to its population centroid. To overcome this issue, we use normalization that captures the distance to the mean in units of standard deviations, as in [Liao et al., 2009] and [Patterson et al., 2006]. In a Euclidean space, the *Mahalanobis* distance is the distance of a test point to the centroid divided by the width of the ellipsoid in the direction of the test point ([Hastie et al., 2009]), given by a variance-covariance matrix Σ. In terms of the distance of a binary vector *x* to mean vector *μ*,

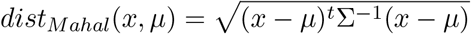

Similarly, we define a corresponding *ASD-Mahalanobis* (ASD-M) distance,

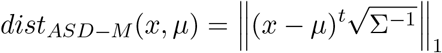

The assumption of *linkage equilibrium* implies a diagonal covariance matrix (comprising only variances) so that the distance of *x_k_* to *M*_1_ for our two metrics is,

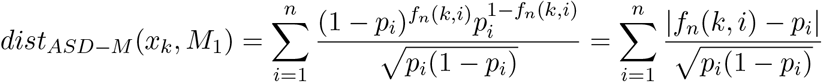

such that our classifier function *D_k_* is

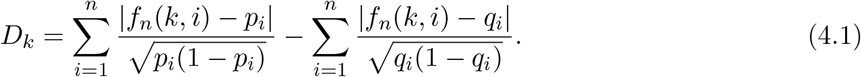

Finally, we would like to incorporate the class priors into our classifier function *D_k_*. Since priors do not naturally enter a distance classifier as they do a Bayesian posterior, we approach this problem by reformulating the expression of *D_k_* for the Bayes and NC classifiers in a similar fashion. The Bayes classifier from Eq.(3.5) can be reformulated as,

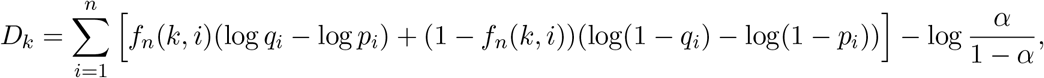

whereas the NC classifier from Eq. (4.1) as,

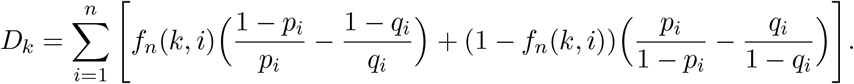

From this we derive the function that best reflects the effect of the prior as −log .

Interestingly, this alternative formulation of *D_k_* provides insight into why different classifiers deviate from the optimal Bayesian, at least for multidimensional binary distributions; in the case of the NC classifier, this is due to 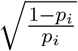 being only a rough *approximation* of − log *p_i_*.

Finally, the we transform the resulting error rate to an expression for *C_n_* as in Eq. (3.4),

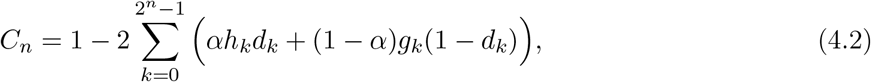

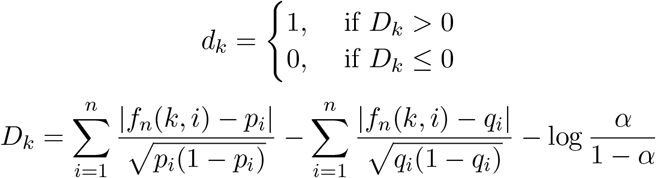

where the genotype frequencies are as in Eq. (2.1).

Nevertheless, despite the use of a normalized metric in the NC classifier, the distance-based *C_n_* does not comply with all required criteria (see Appendix D). We briefly remark here that this deficiency is most probably a characteristic of many other non-optimal classifiers. For instance, we have examined a simple information-theoretic classifier based on the notion of *typical sequences* from [Tal et al., 2017]. Using a closed-form formulation of the error rate of the *cross-entropy classifier* (their Appendix A.2) and allowing for class priors, it is easy to show by counter example that at least two of our criteria are not met: the *performance* criterion (e.g. *p_i_* = 0.1/*q_i_* = 0.3 fails at *n* = 5) and the neutrality criterion (e.g. *p_i_* = 0.03/*q_i_* = 0.2, *n* : 1… 6 and *p_i_* = 0.1/*q_i_* = 0.1 for *n* > 6 fails at *n* = 11).

## 5 Sampling Effects

Our information measure has been defined parametrically, from the underlying properties of genotype distributions across two populations. In practice, however, researchers employ estimates made from sampled data rather than parametric allele frequencies. Simulations of a variety of classification methods on genetic data show that performance is always degraded with smaller population samples, most notably given low values of differentiation ([Cornuet et al., 1999]; [Rosenberg, 2005]; [Tal, 2012a]). Nevertheless, there is information available for classification under these more restrictive circumstances, which we would like to quantify.

The *test error*, also referred to as *generalization error*, is the prediction error over an independent test sample. Here the training set is fixed, and test error refers to the error for this specific training set. A related quantity is the expected prediction error (or expected test error). In practice, estimation of test error conditional on a particular training set is in general difficult, given just the data from that training set. Instead, cross-validation and related methods may provide reasonable estimates of the expected test error ([Hastie et al., 2009]). We therefore extend the information measure to encompass these limiting circumstances.

Denote by *C_n,m_* the *informativeness* under a sampling scenario, effectively an *expectation over all training samples of size m*, where *m* = {*m*_1_,*m*_2_} indicates sample sizes from the two populations. The introduction of this sampling framework calls for a new set of criteria, which are formulated in reference to *C_n_* to replace the existing criteria.

[1*]*Sampling-Effect*: For any sample size m, *C_n,m_* ≤ *C_n_*. This requirement is justified by established observations and theoretical results on reduced classification performance with smaller samples (e.g., [Cornuet et al., 1999]; [Rosenberg, 2005]).
[2*]*Sampling-Convergence:* As sample size m increases *C_n,m_* → *C_n_*. This requirement is justified by common observations on the effect of increasing sample size ([Cornuet et al., 1999]; [Rosenberg, 2005]), also motivated from the *efficiency* quality of any estimator – approaching the underlying parameter being estimated with a probability approaching one as the sample size becomes large.

The introduction of sampling effects requires a reformulation of *C_n_* that distinguishes frequency *estimates* from the *parametric* frequencies. Essentially, the former are used in the expression for the classifier, while the latter are used in the expectation of the conditional error and thus we express a classifier module that is separate from the averaging process. Instead of the misclassification rate, we require a *conditional test error* - the expected error of a classifier over an independent training sample, and the expected test error or the *generalization error* - an average over all training samples of size *m* ([Hastie et al., 2009]). Our goal is to express the *expected test error* for the Bayesian and *nearest-centroid* classifiers in closed form.

Let *m*_1_ and *m*_2_ denote given samples size from the two haploid biallelic populations. Let 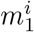 and 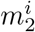 be the average number of allele ‘*A*’ in locus i over all training samples of size *m*_1_ and *m*_2_ respectively. The *maximum likelihood estimator* (MLE) of the true allele frequencies is an unbiased estimator,^4^

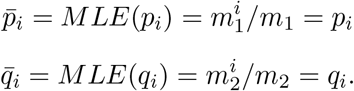

The distributions of the sample mean (allele frequency estimates) *X_i_* and *Y_i_* for populations *P* and *Q* respectively, are binomial,^5^

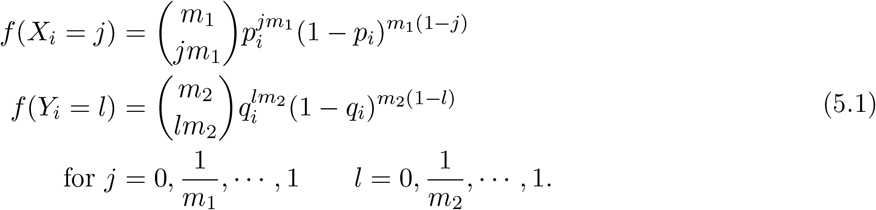

The *expected test error under all training samples of size m* = {*m*_1_, *m*_2_} is the expectation with respect to the sampling distribution of the allele frequencies of the conditional test error *C_n_*(*X, Y*), which is conditional on a particular sample of size *m* of frequency values *X* and *Y*,

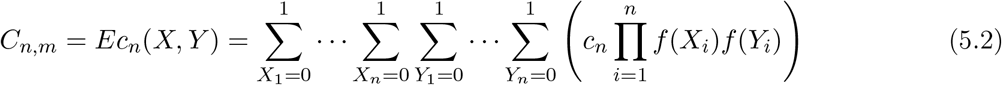

where *c_n_* follows the general formulation in Eq. (3.4),

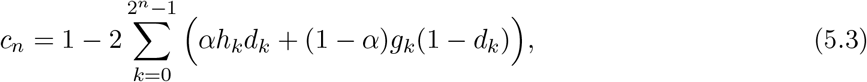

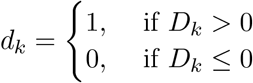

And where the classifier compares genotype *sample* probabilities given a particular sample, following the *Bayes* classifier formulation of Eq. (3.5),

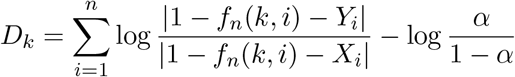

where *h_k_,g_k_,f_n_*(*k,i*) are defined in Eq. (2.1).

*C_n,m_* indeed complies with the two new criteria (see Appendix E for proof).

## 6 Discussion

Much of the work on elucidating factors contributing to the *informativeness* of genetic markers for ancestry inference had focused only on a small number of core factors, namely, higher *informativeness* with additional loci, with wider source population divergence, and with a larger population sample size in the learning phase ([Estoup and Angers, 1998]; [Cornuet et al., 1999]; [Edwards, 2003]; [Rosenberg et al., 2003]; [Wang, 2006]; [Witherspoon et al., 2007]; [Ding et al., 2011]; [Tal, 2012a]; [Tal, 2012b]). A central goal of this paper was to provide a formal grounding for an effective decision-theoretic measure compliant with an extended set of such factors framed axiomatically at the outset, subsequently revealing a host of novel intrinsic properties of both theoretical importance and practical relevance. Although one normally proceeds by explicitly deriving a measure from a set of axioms, we were constrained by the sheer cardinality of this set, adopting a top-down approach instead: demonstrating that a prospective *informativeness* measure adheres to all criteria. Naturally, in any axiomatic framework one strives to include some minimal set of consistent and independent criteria, such that no criterion contradicts nor logically follows from a combination of others ([Rodin, 2014]). The consistency of our framework follows in retrospect from the demonstration of at least one candidate in full compliance (the Bayes-based *C_n_*). The independence aspect is harder to make formal but should be sufficiently intuitively captured from the basic formulations.

### The information-theoretic measure

*Mutual Information* based measures have been widely used in both *feature selection* ([Battiti, 1994]; [Last et al., 2001]; [Grall-Maes and Beauseroy, 2002]; [Huang and Chow, 2005]; [Huang and Rong, 2009]), most effectively exemplified in the *Max-Dependency principle* ([Peng et al., 2005]), and in deriving *ancestry informativeness measures* ([Rosenberg et al., 2003]). The information-theoretic approach has origins more generally in the *infomax principle*, where a subset of features *Y_θ_* should be chosen so that the mutual information between the features and the class label *X, I*(*Y_θ_;X*), is maximized (equivalently, that the conditional entropy *H*(*X*|*Y_θ_*) is minimized). Some formal justification for the relevance of this principle to classification tasks is given by *Fano’s inequality*, which relates the conditional entropy between the source and destination in a communication channel to the transmission error ([Zhao et al., 2013]). As a result, mutual information can be used to express both an upper and lower bound for the Bayes error rate ([Brown et al., 2012]).

Nevertheless, much of the justification for employing *mutual information* in *ancestry informativeness* or *feature selection* is simply based on its usefulness in practice. For instance, [Peng et al., 2005] utilize a multivariate mutual information for feature selection since “minimal error usually requires the maximal statistical dependency of the target class c on the data distribution in the subspace Rm (and vice versa)”. This approach is effectively epitomized by a recent analysis of filters for feature selection, where the authors explain that “an intuitive [filter] J would be some measure of correlation between the feature and the class label-the intuition being that a stronger correlation between these should imply a greater predictive ability when using the feature” ([Brown et al., 2012]).

Here we have shown that *multilocus ancestry informativeness* measures based on *mutual information* that also account for population priors (such as the *multilocus version* of *I_n_* proposed in [Rosenberg et al., 2003]) do not properly incorporate the information available from such priors. In the context of our axiomatic framework, this amounts to non-compliance with the *priors* criterion, which requires any measure of *informativeness* to approach its maximum when priors approach their extremes (reflecting a high discrepancy in source population sizes). This is of significance since *informativeness* ultimately represents the *value* of information given by the data (loci and priors) that an effective classifier can utilize in making correct inferences. In essence, it is a proxy for the performance of effective classifiers in a given setting, the level of certainty they can achieve. Indeed, we have shown that the decision and information theoretic measures may give different relative rankings to SNP panels under certain values of the priors. This ranking discrepancy is incongruent with the claim in [Rosenberg et al., 2003] that ORCA (analogous to our Bayes-based *C_n_*) and *I_n_* are highly correlated and produce similar estimates of panel ranking so that one ‘can proceed using only one statistic’. This is most likely a consequence of our focus here on multilocus rather than single-locus *informativeness*, along with the incorporation of population priors. Indeed, numerical simulations indicate that the ranking discrepancy between *C_n_* and *I_n_* is accentuated with a larger panel size (n) and with greater inequality in population priors – a rigorous analysis of which is not further pursued here and is left for future work.

### Properties of *C_n_*

We have revealed properties of the Bayes-based *C_n_* that broadly pertain to two important aspects of this measure: subadditivity across loci and populations, and an interplay of information between the prior and the genetic variants. The *loci-subadditivity* property, which compares the sum of *informativeness* of two genetic sequences to that of the aggregated sequence, can be viewed as a generalization of a theorem by [Gattepaille and Jakobsson, 2012] on the *informativeness* of two-locus haplotypes of biallelic markers *in the absence of linkage disequilibrium within populations*. These authors were concerned with the effect of LD on the strength and sign of the *gain of informativeness for assignment* (GIA), a term they used to describe as similar inequality as above, utilizing the mutual information based *I_n_* of [Rosenberg et al., 2003], concluding that “although there are a number of predictable behaviors of GIA-such as that GIA ≤ 0 when markers are in linkage equilibrium and that GIA is often large for cases where private alleles exist–GIA is not a trivial function of LD or allele frequencies”. Their primary motivation was to use this statistic as a heuristic for deciding whether to combine SNPs into haplotypes for improved accuracy in population assignment schemes, specifically targeting that part of LD that is due to physical linkage in the genomes, i.e., to handle the issue of having markers close along a chromosome (a type of data is typically pruned before further use in assignment schemes).

The *population-subadditivity* property allows interpreting *C_n_* as a distance *metric* between populations *relative to any given number of loci*. By definition, a metric on a set *X* is a *distance function d* : *X* × *X* → ℝ that satisfies four conditions: *d*(*x,y*) ≥ 0 (the non-negativity of *informativeness*), *d*(*x,y*) = 0 if and only if *x* = *y* (the zero criterion), *d*(*x, y*) = *d*(*y,x*) (the *invariances* criterion for class symmetry) and *d*(*x,z*) ≤ *d*(*x, y*) + *d*(*y,z*) (this *population-subadditivity* property). This feature of *C_n_* is in stark contrast to standard differentiation measures such as *F_ST_ which are not metrics* (see [Tal, 2013] for analysis of several population genetic distances).

The *prior-washout* property describes the diminished effect of the population prior as additional loci are included in *C_n_*. Although this phenomenon is a direct consequence of the *performance* criterion, we have chosen to separately highlight it since it offers another perspective on the balance between the informative roles of the population prior vs. the genetic variants. For Bayesian classifiers, the washout effect also emerges from taking an information-theoretic perspective, by utilizing the *asymptotic equipartition property* (AEP) of relative entropy typical sets ([Cover and Thomas, 2006, p. 388]).

The interplay between the effects of the population prior vs. that of the genetic variants on our *C_n_* is most sharply exemplified by the *prior-sensitivity*, *prior-dominance and uninformative-loci* properties. We have shown that while *C_n_* is always sensitive to fluctuations in the prior, there are cases (which we characterize in detail in the proofs) where an additional locus is uninformative (i.e., *C_n_* is invariant) given any value of the prior; and more surprisingly, we show that for any given set of loci, prior extremities beyond certain thresholds (determined by allele frequencies at these loci, Eqs. (C.2), (C.3) in Appendix C) render the information from the loci *completely redundant for the assignment task*. At this range of high disparity of population size, *informativeness* is only a function of the priors, with the consequence that effective classifiers may then simply assign unknown genotypes to the largest source population (as evidenced by the prior and likelihood terms of the log-posterior ratio of a Bayes classifier).

### Feature Selection

Searching for the best *m* features out of the *n* available for the classification task is known to be a NP-*hard* problem and thus exhaustive evaluation of possible feature subsets is usually unfeasible in practice due to the large amount of computational effort required ([Huang and Rong, 2009]). This is why single-locus rather than multilocus based heuristics are common ([Rosenberg, 2005]). While *C_n_* is not intended as a metric or heuristic for implementing feature selection schemes as it is inherently multilocus in nature, it could still be employed for related tasks. Primarily, it can be used for comparing between a small number of given SNP panels, for verification subsequent to a standard feature selection task. Moreover, the *delta* criterion implies the possibility of ranking single loci for their contribution to an already existing panel in the particular case where a subset of loci has completely overlapping population frequency differences. We emphasize in this context that it is generally not possible to rank loci for their contribution to total informativeness, due to their interdependence and redundancies. Indeed, it has long been recognized in feature selection that combinations of individually good features do not necessarily lead to good classification performance. In other words, selecting the loci that are individually most informative does not necessarily produce the optimal panel, or more compactly put, “the m best features are not the best m features” ([Peng et al., 2005]).

In the context of feature selection, it is common to make a distinction between filter and wrapper methods. Filter type methods select variables regardless of the classification model, and are based only on general principles like a correlation of features with the predicted variable. Such methods are particularly effective in computation time for a large set of features, and in robustness to overfitting when the number of observations is small. In contrast, a wrapper is a feature selector that convolves with a classifier (e.g., naive Bayes classifier), with the direct goal of minimizing the classification error of that particular classifier. Usually, wrappers can yield high classification accuracy for a particular classifier at the cost of high computational complexity and less generalization of the selected features on other classifiers.

While *C_n_* has features of both filters and wrappers, it is more appropriately categorized as a wrapper, since it incorporates multiple features and has an explicit relation to the classification model (essentially derived from a classifier error rate). In this context is worth noting that our Bayes-error based *C_n_* is not prone to *overfitting* since it effectively uses an infinite training sample (i.e., it is parametrized by underlying allele frequencies). Moreover, the finite-sample *C_n,m_* also does not suffer from *overfitting* as it employs estimates of single SNP frequencies, rather than multilocus genotype frequencies.

### Estimation from finite samples

Previous work on incorporating finite population samples in ancestry *informativeness* measures and in *panel selection* had proceeded by simply replaced true allele frequencies with sample frequencies in the *informativeness* measure. This approach, however, cannot provide general insight to the diminished informational content due to working with frequency estimates (e.g., [Rosenberg et al., 2003] in their Bayes-based ORCA and *mutual information*-based In for both the single- and multilocus formulations; see [Sampson et al., 2011] for a discussion of this issue). While this approach is may be appropriate in the context of practical feature selection schemes based on particular training samples, it does not convey the intrinsic degrading effect on *informativeness* resulting from finite samples. Closer to our approach is the analysis in [Rosenberg, 2005] of a *performance function* for panel selection, which proves (Theorem 7) a result akin to our *sampling-convergence* criterion of *C_n,m_* for the Bayes-based ORCA statistic computed by simulation using sample allele frequencies. However, this result only captures the *conditional test error*, whereas we target the *expected* test error – an expectation of the former over all possible samples of size *m*. That treatment also includes a performance result characterizing the effect of additional loci (Corollary 8 of Theorem 7) given some minimal sample size threshold – an analysis which we do not undertake here for *C_n,m_*.

### Distance-based Methods

We have also analyze a simple distance-based classifier and derived a model of its error rate to see whether other effective classifiers can serve as a basis for *informativeness*. The construction of a nearest-centroid classifier for the high-dimensional discrete genetic data has revealed that simple genetic metrics such as the *allele sharing distance* do not take into account the variance-covariance matrices of the distribution of genotypes of each population (theoretically, if the centroids lie on *opposite* vertices of the hypercube genotype space, the classifier may associate up to 2(*n* − 2) vertices to the ‘wrong’ centroid). Such metrics effectively ignore the width of the distributions in the direction of individual to population-mean comparison. This issue probably explains the weak performance of the misclassification rate *C_c_* from Fig. 2 in [Witherspoon et al., 2007], with data from low MAF sites (“rare” alleles, *MAF* < 0.1). Nevertheless, other treatments have ignored this issue, e.g., [Degen et al., 2017] who use a k-Nearest Neighbor (*k*-NN) algorithm for classification to one of a collection of source populations based on simple pairwise individual allele-sharing distance.

To overcome this issue, we developed a normalization akin to a *Mahalanobis* distance adopted to the discrete nature of the genetic space. In a Euclidean space, the *Mahalanobis* distance is the distance of a test point to the centroid divided by the width of the ellipsoid in the direction of the test point ([Hastie et al., 2009]) and is normally applied within classification techniques such as the k-Nearest Neighbor (k-NN) or linear and quadratic discriminant analysis (LDA, QDA). The use of a *Mahalanobis* distance is common procedure in classification schemes involving *continuous gene expression* data. [Dudoit et al., 2002] have also implemented similar normalization for nearest-centroid classification of gene expression profiles, using *Diagonal linear discriminant analysis* (DLDA). However, their model is different in a few crucial respects from our model: they assume normality of the underlying data distributions and equal class densities. A similar approach was adopted in [Tibshirani et al., 2002, Tibshirani et al., 2003] describing a method of classifying gene expression test samples according to *closest shrunken centroids*, also standardizing by variance estimates.

An approach that bears a stronger resemblance to our SNP distribution model is described in [Patterson et al., 2006]. The paper describes an application of PCA for genetic biallelic data using standardized distances. Each entry in the PCA matrix is normalized by the variance of *C*(*i, j*), the number of occurrences of the MAF allele for locus *j* from individual *i*. The authors report that “We verified (unpublished data) that the normalization improves results when using simulated genetic data, and that on real data known structure becomes clearer.” Even closer to our distance-approach is the work of [Liao et al., 2009] which introduces a classifier for SNP data genotypes with distance normalization by variance and a similar correction for population priors via a logarithmic term. However, since it employs a *shrinkage* centroid deriving a simple expression for its error rate would be hardly feasible.

The naïve Bayes classifier upon which *C_n_* is formulated derives multilocus genotype frequencies under an implicit assumption of linkage equilibrium (LD). Indeed, we would not normally expect to see linkage disequilibrium within distinct populations, except between markers that are in close proximity (physical linkage, pruned in common preprocessing procedures) or in cases of recent admixture ([Pritchard et al., 2000]). At a population level, admixture manifests as LD between markers reflecting a shared history of descent, and invalidating the independence assumption. Haplotype based analysis was devised to potentially harness this information ([Lawson et al., 2012]). A possible approach for extending our model for *ancestry informativeness* to take advantage of information from LD that exists in structured populations, might involve including variance-covariance matrices that specify pairwise LD estimates for normalization in an NC classifier, instead of the diagonal covariance matrices. This extension is left for future work.

### Conclusion

In this paper we have taken an axiomatic approach to select a measure for *ancestry informativeness*. We have shown that a measure based on an optimal Bayes classifier complies with the full set of proposed criteria, in contrast to a popular measure based on mutual information. The core deficiency of the information theoretic approach is in properly incorporating the information available from population priors. This is exemplified in decision and information theoretic measures providing different relative rankings to SNP panels under certain population priors. The measure based on a Bayes error was moreover shown to possess several surprising properties which characterize the interplay between information originating from the priors and the genetic markers. We have also analyzed a distance-based classifier adopted to discrete high-dimensional population genetic data, as an instance of an effective classifier which nevertheless cannot form a basis for satisfactory multilocus ancestry *informativeness* measure. Finally, we have extended the framework to formally quantify the inherent degrading effect on *informativeness* from using finite population samples in the learning phase.

## Acknowledgements

Special thanks to Jim Portegies and Guido Montúfar for fruitful technical discussions. We also acknowledge Jürgen Jost and the Max Planck Institute for Mathematics in the Sciences for its generous support and the platform to present an earlier form of these ideas in an internal seminar.

## Appendix A The relation between the Bayes error and some generalization of the variational distance

### A.1 The relation

Here we develop a proof for the general case of *unequal priors*. It is required to show that,

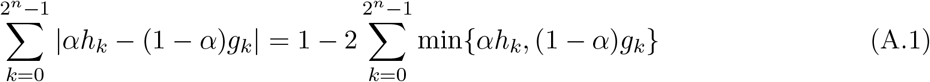

where,

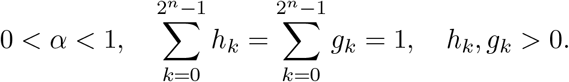

#### Proof.

In fact, we note that |*x* − *y*| = *x* + *y* − 2 min{*x, y*} for all *x, y*. Therefore

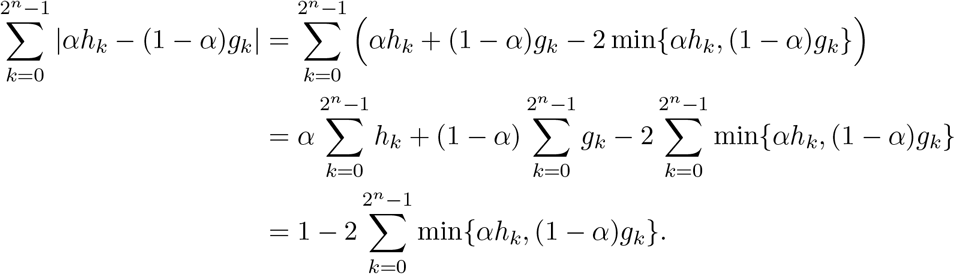

### A.2 Other representations of *C_n_*

Reminder:

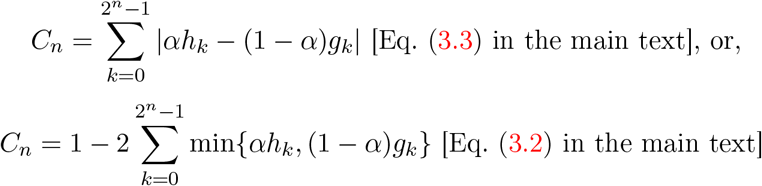

where,

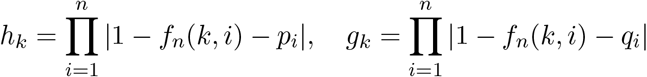

and *f_n_*(*k, i*) is the *i^th^* bit of *k* in its binary form.

To simplify the proofs, we also use another representation of *C_n_*

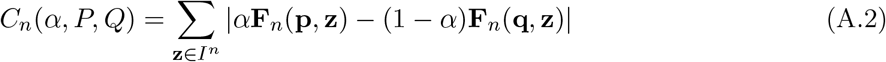

where *I^n^* = {0,1}^*n*^ and 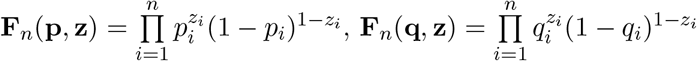.

#### Proposition 1.

*The Eq*. (A.2) *is equivalent to the Eq*. (3.3).

*Proof*. In fact, we have an isomorphism between each *k* = 0, … , 2^*n*^ − 1 and one corresponding element 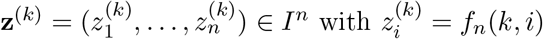. Moreover, we have

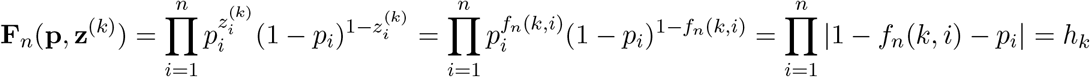

and similarly **F**_*n*_(**q, z**^(*k*)^) = *g_k_*. This implies the proof.

#### Remark 1.

*From Eq*. (A.1) *and Eq*. (A.2) *imply*

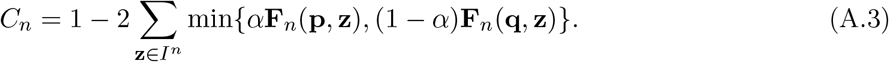

To simplify the proofs, we will use all four these representations of *C_n_* (i.e. (3.2), (3.3), (A.2), (A.3)).

## Appendix B

Proof of compliance of Bayes *C_n_* with all criteria

1. *Zero*: (⇐): If *P* = *Q* (i.e. *p_i_* = *q_i_*, ∀*i* = 1, … , *n*) and *α* = 0.5 then we have **F**_*n*_(**p, z**) = **F**_*n*_(**q, z**) for all **z** ∈ *I^n^*. Thus from Eq. (A.2), and since the sum of all genotype frequencies from a single population is 1,

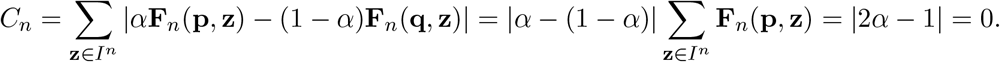

(⇒): Conversely, we need to show that *C_n_* = 0 implies *P* = *Q* (i.e., *p_i_* = *q_i_*, ∀*i* = 1, … ,*n*) and *α* = 0.5. First, notice that if *C_n_* = 0 then trivially each summand of Eq. (A.2) must be zero, i.e. *α***F**_*n*_(**p, z**) = (1 − *α*)**F**_*n*_(**q, z**), ∀**z** ∈ *I^n^*. By summing for all **z** ∈ *I^n^* we have *α* = 1 − *α* or *α* = 0.5. Therefore **F**_*n*_(**p, z**) = **F**_*n*_(**q, z**), ∀**z** ∈ *I^n^*. Denote by 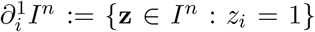. Then we have for every *i* = 1, … , *n*

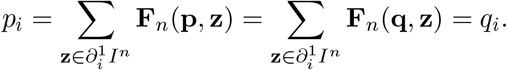
2. *Performance*: See proof in [Tal, 2012b, Appendix B.1], and Fig. B.1 for illustration.
3. *Asymptotics*: See proof in [Tal, 2012b, Appendix B.2], and Fig. B.2 for illustration.
4. *Neutrality*: We need to prove that *C_n_* = *C*_*n*+1_ if *p*_*n*+1_ = *q*_*n*+1_. In fact, by denoting 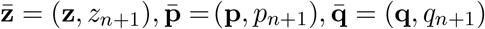, we have

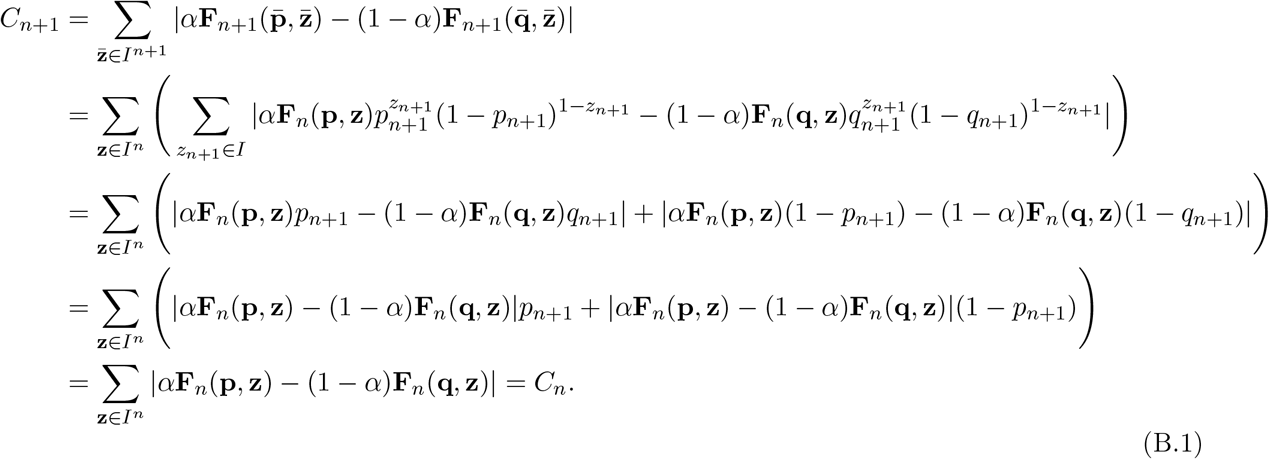
5. *Continuity*: From Eq. (A.2), *C_n_* is a sum of absolute values of continuous functions and therefore has no singularities, and since *p_i_,q_i_* and *α* are real-valued parameters, *C_n_* is continuous with respect to its parameters.
6. *Dominance*: (⇒) We need to prove that for a fixed *n* and some fixed *α* ∈ (0, 1), if there exists *i* ∈ {1, … , *n*} such that |*q_i_* − *p_i_*| → 1 then *C_n_* → 1. In fact, without loss of generality we assume that *p_n_* → 0 and *q_n_* → 1. We note that

- If *z_n_* = 0 then 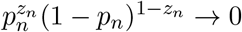 and then **F**_*n*_(**p, z**) → 0
- If *z_n_* = 0 then 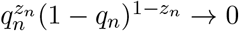 and then **F**_*n*_(**q, z**) → 0. Therefore min{*α***F**_*n*_(**p, z**), (1 − *α*)**F**_*n*_(**q, z**)} → 0 for all **z** ∈ *I^n^*. This results in *E_n_* → 0 and consequently *C_n_* → 1.

**Fig. B.1:**
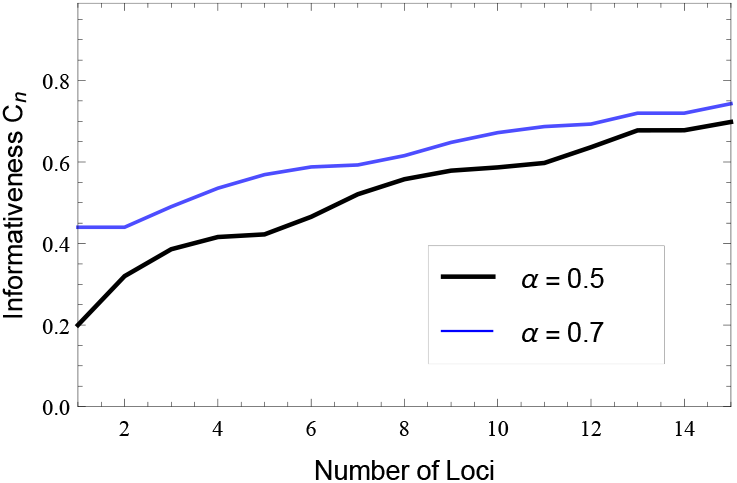
*C_n_* based on Bayes error complies with *performance*. Shown are two cases corresponding to different class priors, with allele frequencies *p*_1–12_ = 0.1/*q*_1–12_ = 0.3,*p*_13_ = 0.05/*q*_13_ = 0.3,*p_14_* = 0.24/*q*_14_ = 0.2,*p*_15_ = 0.01/*q*_15_ = 0.15.

**Fig. B.2:**
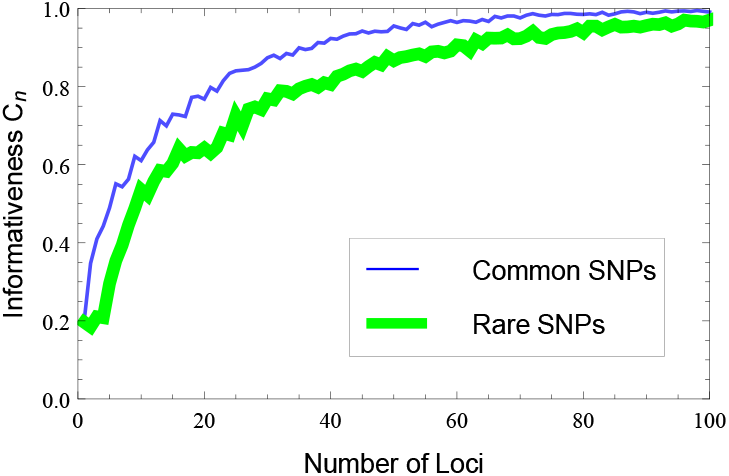
*C_n_* based on Bayes error complies with *asymptotics*. Two cases are shown (using Monte Carlo simulations), common (*p*_1_ = 0.2/*q*_1_ = 0.45) and rare (*p*_1_ = 0.02/*q*_1_ = 0.12) SNPs and prior *α* = 0.4.

(⇐) Conversely, assume that *C_n_* → 1 for a fixed *n* and some fixed *α* ∈ (0,1), we need to prove that there exists *i* ∈ {1, … , *n*} such that |*q_i_* − *p_i_*| → 1. In fact, we first note that **p** ≠ **q** because otherwise |2*α* − 1| = *C_n_* → 1 which is a contradiction. Moreover, because *C_n_* satisfy the neutrality and invariant property, we can assume that *p_i_* < *q_i_* for all *i*. Now, put 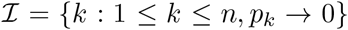 and 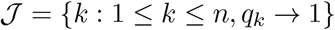. Because *C_n_* → 1, it implies

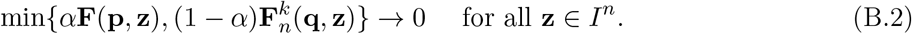

By choosing **z** = (0, … , 0) we imply 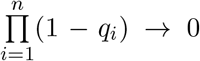, therefore 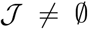. Similarly, by choosing **z** = (1, … , 1) we imply 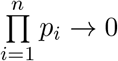, therefore 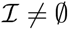. Assume that In 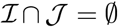 then we can choose **z** ∈ *I^n^* such that

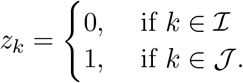

Note that

- 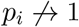 for 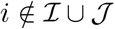 (otherwise, it implies *q_i_*(> *p_i_*) → 1 and contradicts to the definition of 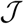);
- 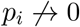 for 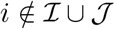 and 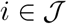 (from the definition of 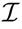).

Therefore we have

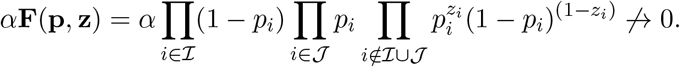

Similarly, we have 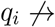 for 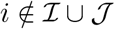; 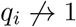 for 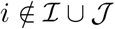 and 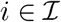 and therefore

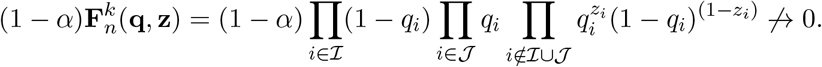

It implies that 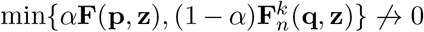 which contradicts to (G.4). Thus 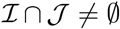 which implies the proof (see Fig. B.3 for illustration).

7. *Delta*: For simplicity in notation, we examine *n* + 1 loci (where *n* can also be 0), and the proof proceeds without loss of generality with respect to locus *n* + 1 with allele frequencies *p* and *q*. First, express *C*_*n*+1_ as in Eq. (B.1),

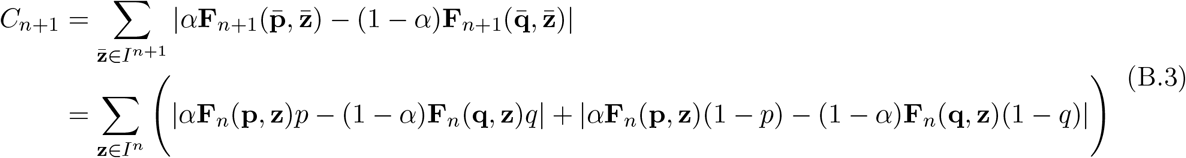 Each pair of *absolute-value* terms can be seen as a sum of two *V*-shaped functions of *q*, without loss of generality (illustrated in Fig. B.4). Let us examine the first pair, indexed by **z**. The zero ‘tip’ of one *V*-shaped function occurs where,

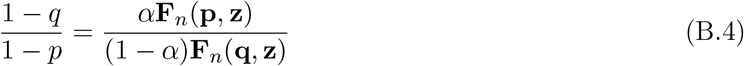

while the zero tip of the other occurs where,

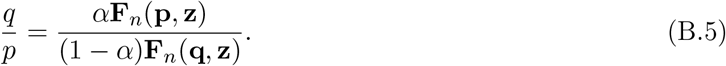 Assume, without loss of generality, that *q* > *p* at the zero-tip described by Eq. (B.4). Since the RHS equals the RHS of Eq. (B.5), then *q* < *p* at the zero-tip described by Eq. (B.5). Now, since the absolute values of the slopes of all *V*-shaped function pairs are equal, their sum is a 3-piece-wise linear convex function where the center section is of slope zero (Fig. B.4). This is also the case for any paired sum in Eq. (B.3), and since the sum of convex functions is convex, *C*_*n*+1_ is necessarily convex. Crucially, the slope-zero sections of all these 2^*n*^ convex functions (as a function of *q*) necessarily partially overlap, so that their sum *C*_*n*+1_ also has a slope-zero section (Fig. B.4). Thus, if *q* < *p* then substituting a lower value for *q* can only increase *C*_*n*+1_; similarly, if *q* > *p* then substituting a higher value for *q* can only increase *C*_*n*+1_.
8. *Invariances*: [a] since the genotype probabilities **F**_*n*_(**p, z**) and **F**_*n*_(**q, z**) in Eq. (A.2) are each a commutative product of allele frequencies from all loci, *C_n_* is invariant to different ordering of loci; [b] *C_n_*(*α, P, Q*) = *C_n_*(1 − *α, Q, P*) follows from the presence of an absolute value in the formulation of Eq. (A.2); [c] the simultaneous substitution of *p_i_* with (1 − *p_i_*) and *q_i_* with (1 − *q_i_*) simply changes the order of the summation terms in Eq. (A.2) and thus does not affect *C_n_*. We illustrate this with regard to C_2_:

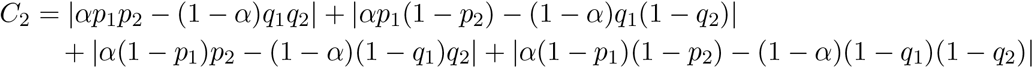 Now, after the substitution (termed here 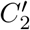) we get,

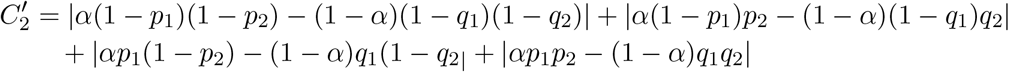
9. *Prior*: If *α* → 0 or *α* → 1 then one of the two terms within the sum in Eq. (A.2) diminishes to zero and what remains in the limit is the sum over all genotype probabilities in one population, which equals 1, therefore *C_n_* → 1.

**Fig. B.3:**
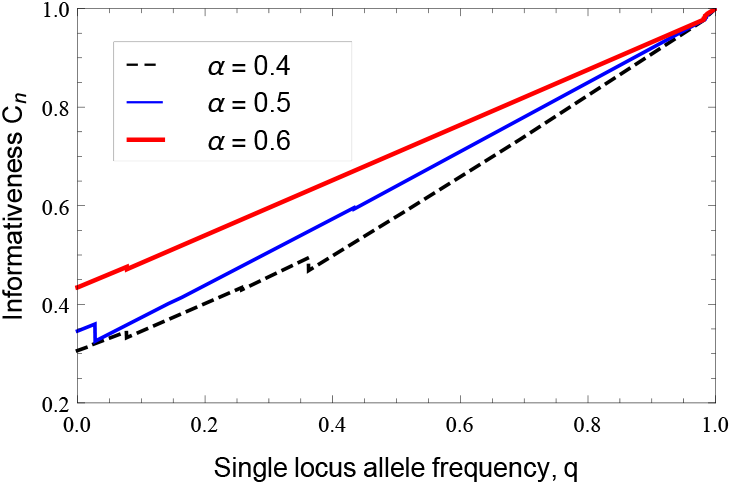
*C_n_* based on Bayes error complies with *Dominance: C_n_* → 1 as |*q_i_* − *p_i_*| → 1. Shown here are three cases corresponding to different class priors, for 5 loci where *q*_5_ changes from ~ 0 through ~ 1 while *p*_5_ ~ 0; the other 4 loci have frequencies: *p*_1_ = 0.01/*q*_1_ = 0.3,*p*_2_ = 0.2/*q*_2_ = 0.35,*p*_3_ = 0.4/*q*_3_ = 0.3,*p*_4_ = 0.1/*q*_4_ = 0.15.

**Fig. B.4:**
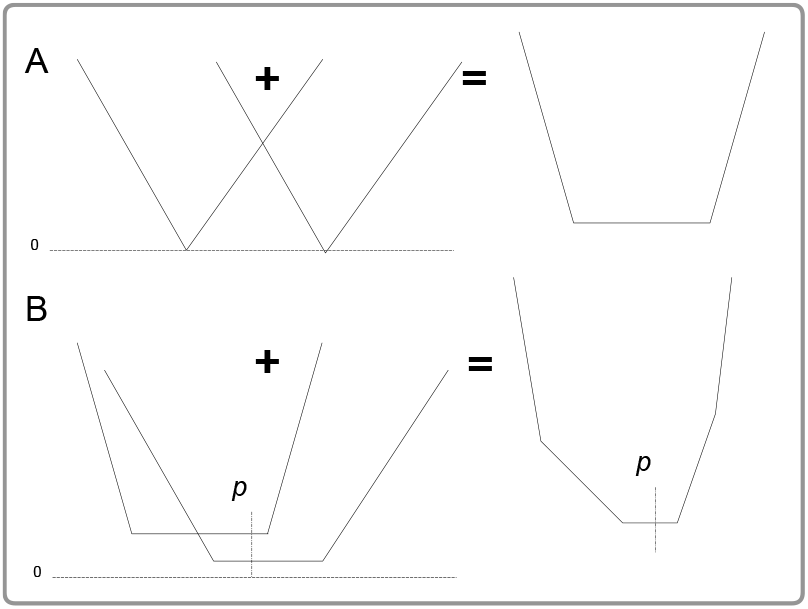
A schematic for the proof of the delta criterion.

## Appendix C Proof of the properties of the Bayes-based *C_n_*

a. *Loci subadditivity (w/out priors):* we need to show that *C_n_+_m_*(*P*║*P*′, *Q*║*Q*′) ≤ *C_n_*(*P, Q*)+*C_m_*(*P*′, *Q*′). Let 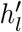 designate the 2^*m*^ genotype frequencies associated with frequency vector *P*′, and similarly 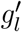 with reference to frequency vector *Q*′. Then from Eq. (3.3) it directly follows that,

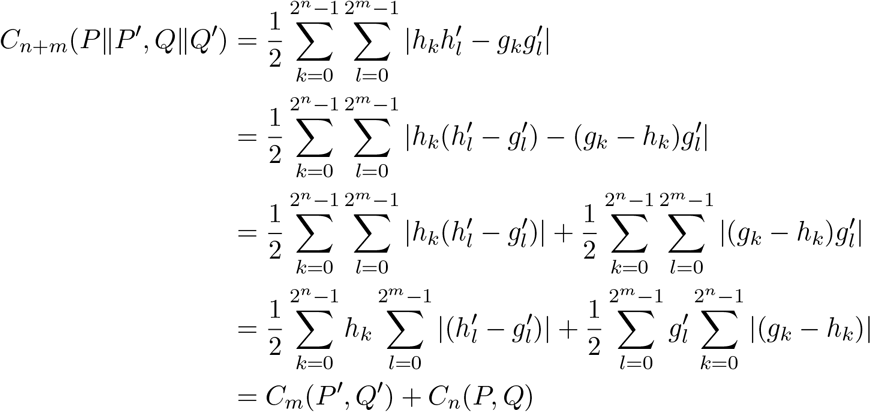

where the transition between lines 2 and 3 is due to the triangle inequality (|*a* − *b*| ≤ |*a*| + |*b*|), and the transition between lines 4 and 5 is due the sum over all genotype frequencies at any population equaling 1. We note that when general priors are incorporated in *C_n_*, this property may fail. We may nevertheless allow for priors by introducing a correction term |1 − 2α|,

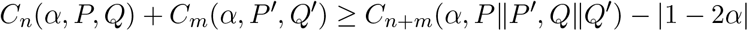 In fact, we have

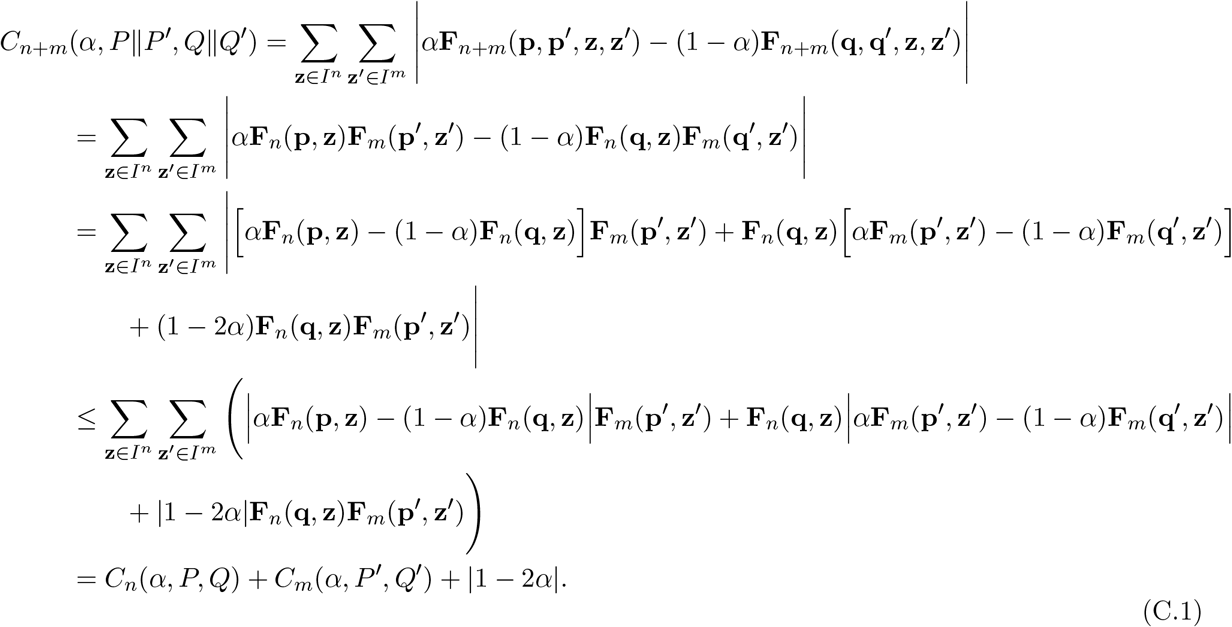
b. *Population-subadditivity (w/out priors):* the compliance of *C_n_*(*P, Q*) with the triangle inequality follows from a formulation in the form of the variational distance, as per Eq. (3.3). [Khosravifard et al., 2007] prove that such distance indeed satisfies the triangle inequality.
c. *Prior-washout:* This property is a direct implication of the *performance* criterion, since the minimum with respect to α also increases, *C*_*n*+1_(*α*) ≥ *C_n_*(*α*) (see Fig. 5), and asymptotically follows from the *asymptotics* criterion *C_n_*(*α*) → 1 with *n*, such the prior is completely washed out at the limit. This property can also be inferred from an information-theoretic perspective on a Bayes classifier ([Cover and Thomas, 2006, p. 388]). The likelihood ratio with priors *π*_1_ and *π*_2_,

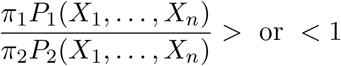

can be formulated as,

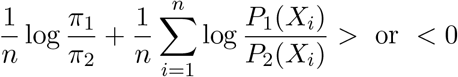

where from the AEP the second term tends to *D*(*P*_1_║*P*_2_) or −*D*(*P*_2_║*P*_1_) in accordance to *P*_1_ or *P*_2_ being the true source distribution, while the first term tends to 0, thus the effect of the prior washes out.
d. *Prior-sensitivity:* We first show that *C_n_* as a function of the prior *α* is convex. Notice from Eq. (3.3) that it is a sum of *V*-shaped convex functions of *α, f*(*α*) = |(*h_k_* + *g_k_*)*α* − *g_k_*| and therefore is itself a convex function. To find *α* giving the minimum of *C_n_*(*α*) notice that the minimum must occur at one or more of the 2^*n*^ ‘singularity’ points of this piecewise linear convex function, given simply by,

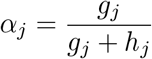

and therefore, the minimum occurs at,

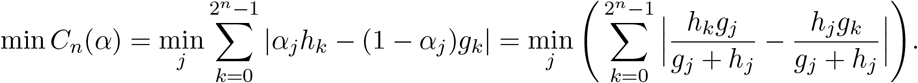 To show that *C_n_* is never invariant to fluctuations in *α*, i.e., that this minimum is unique, we need to prove there are no domains where the slope of *C_n_*(*α*) is zero. The slope of each linear section of *C_n_*(*α*) is some combination of ±(*h_k_* + *g_k_*), the coefficient of *α*. For the slope to be zero this sum over all genotypes has to be exactly zero, a situation which is almost surely (with probability 1) impossible since the allele frequencies are real-valued parameters and thus genotype frequencies never exactly equal between populations.
e. *Prior-dominance:* We need to show that for any combination of allele frequencies there always exist two thresholds strictly in (0,1), 0 < *α*_0_ < *α*_1_ < 1, such that a prior that is more extreme than these thresholds fully determines *C_n_*, i.e., all the genetic loci become effectively *uninformative.* In fact, we note that *α***F**_*n*_(**p, z**) − (1 - *α*)**F**_n_(**q, z**) ≥ 0 if and only if

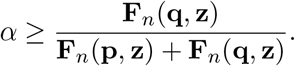 Therefore, by putting

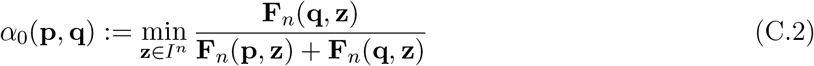

and

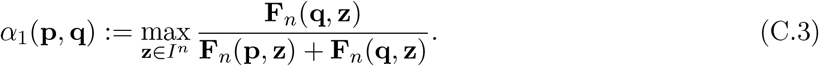 It implies immediately that 0 < *α*_0_(**p, q**) ≤ *α*_1_(**p, q**) < 1. Moreover, for all *α* ≥ *α*_1_(**p, q**) we have

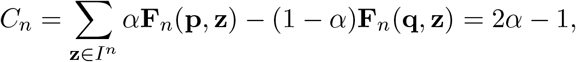

and for all *α* ≤ *α*_0_(**p, q**) we have

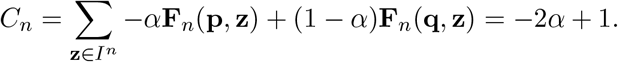 From *C_n_*(*α*_1_) = 2*α*_1_ − 1, *C_n_*(*α*_0_) = 1 − 2*α*_0_ and the *non-negativity* criterion it also follows that 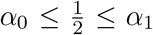. Now, since *α*_1_ and *α*_0_ are continuous functions of *p_i_* and *q_i_* (as **F**_*n*_(**p, z**) and **F**_*n*_(**q, z**) are simply products of those frequencies), slight fluctuations in any allele frequency may induce only a slight change in the thresholds *α*_1_ and *α*_0_, so that the prior remains within the range for which *C_n_* = |2*α* − 1|. More formally, for any *ε* > 0, and prior *α* = *α*_1_ + *ε* or *α* = *α*_0_ − *ε* strictly between 0 and 1, *C_n_* = |2*α* − 1| and is *invariant to slight fluctuations in allele frequencies*. From the *uninformative-loci* property and the *delta* criterion it immediately follows that any locus *i*, for which *C_n_* is invariant to small fluctuations in *p_i_* or *q_i_*, is uninformative; i.e., it does not contribute information in the sense that *C_n_*(*α*) = *C*_*n*−1_(*α*), where this locus is excluded in *C*_*n*−1_. Crucially, this implies that for any prior within the invariant range for *C_n_*(*P, Q*), i.e., *α* ≥ *α*_1_ or *α* ≤ *α*_0_, *all loci* are effectively *uninformative*. To see this, we need to show that the prior thresholds of *C_n_* is included in those of *C*_*n*+1_, i.e. 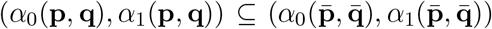; consequently, the exclusion of any m loci would result in *C_n_*(*α*) = *C_n−m_*(*α*) = *C*_0_ = |1 − 2*α*|. In fact, we have from Eq. (C.3)

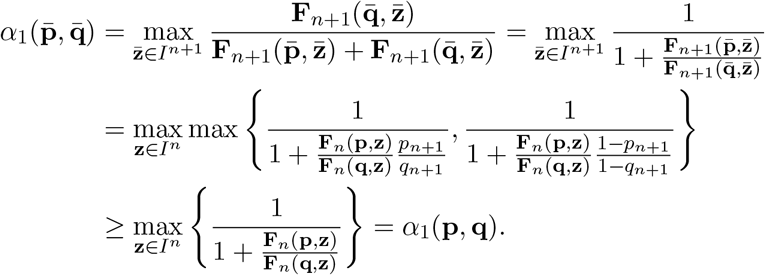 By a similar proof we also have 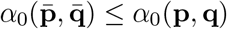. It completes the proof. Note that calculating the thresholds *α*_1_ and *α*_0_ is of exponential complexity *O(k)* = *O*(2^*n*^), but we can achieve a much better result, linear in *n*, i.e., *O*(*n*). This is possible from the following simple transformation,

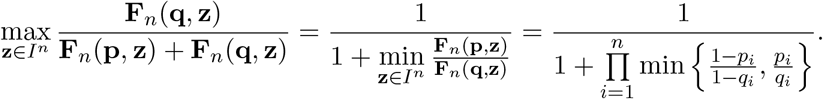
f. *Uninformative-loci:* We need to show that formally for any *P* and *Q* with *n* loci, and without loss of generality,

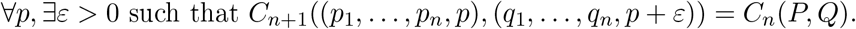

### Proof.

The equality *C_n_* = *C*_*n*+1_ above occurs iff the following inequalities *k* : 0 to 2^*n*^ − 1 from the proof of the performance criterion ([Tal, 2012b, Appendix B.1]) are equalities,

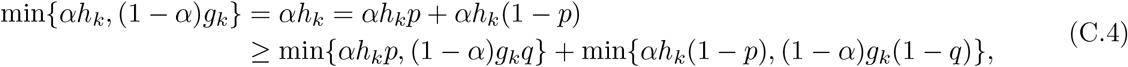

and

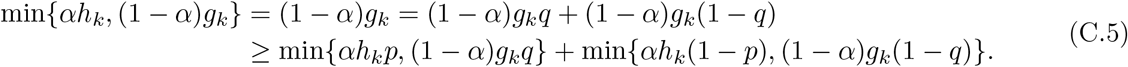

This in turn occurs when, following Eq. (C.4), *αh_k_p* ≤ (1 − *α*)*g_k_q* and *αh_k_*(1 − *p*) ≤ (1 − *α*)*g_k_*(1 − *q*) for each *k* : 0 to 2^*n*^ − 1 for which min{*αh_k_*, (1 − *α*)*g_k_*} = *αh_k_*.

Now write instead of *q, p* + *ε*: *αh_k_p* − (1 − *α*)*g_k_*(*p* + *ε*) and *αh_k_*(1 − *p*) ≤ (1 − *α*)*g_k_*(1 − *p* − *ε*). We get,

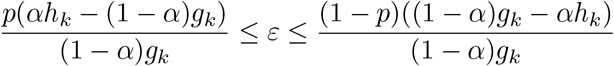

for each *k* : 0 to 2^*n*^ − 1 for which *αh_k_* ≤ (1 − *α*)*g_k_*, and therefore the LHS is non-positive and the RHS is non-negative, and as expected, 0 < *ε* < 1. Also, when following Eq. (C.5), *αh_k_p* ≥ (1 − *α*)*g_k_q* and *αh_k_*(1 − *p*) ≥ (1 − *α*)*g_k_*(1 − *q*) for each *k* : 0 to 2^*n*^ − 1 for which min{*αh_k_*, (1 − *α*)*g_k_*} = (1 − *α*)*g_k_*.

Now write instead of *q,p* + *ε*: *αh_k_p* ≥ (1 − *α*)*g_k_*(*p* + *ε*) and *αh_k_*(1 − *p*) ≥ (1 − *α*)*g_k_*(1 − *p* − *ε*). We get,

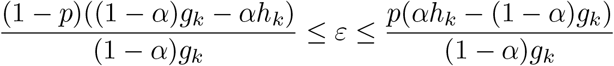

for each *k* : 0 to 2^*n*^ − 1 for which *αh_k_* ≥ (1 − *α*)*g_k_*, and therefore the LHS is non-positive and the RHS is non-negative, and as expected, 0 < *ε* < 1.

Thus unless *alh_k_* = (1 − *α*)*g_k_* for some *k* : 0 to 2^*n*^ − 1, then for any p there is always a non-degenerate range around zero, such that for any *ε* within that range, the inclusion of an extra locus with allele frequencies *p* and *p* + *ε* does not increase *C_n_*.^6^

The range for *ε* is, finally, the overlap of all such ranges (since they are satisfied simultaneously),

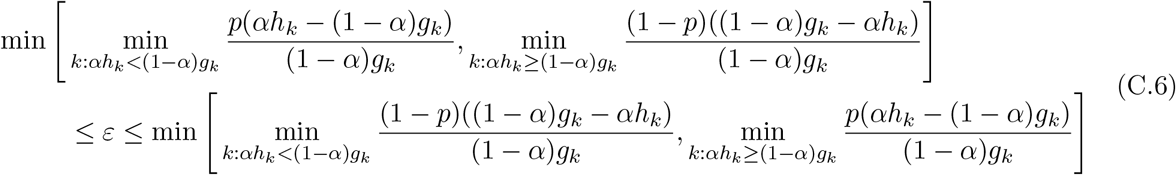

where the internal minimums are taken over *k* : 0 to 2^*n*^ − 1. Since the probability for the occurrence of *αh_k_* = (1 − *α*)*g_k_* for any *k* ranging over all possible genotype frequencies is negligibly small, we can conclude that the theorem is true, *almost surely*.^7^

**Fig. C.1:**
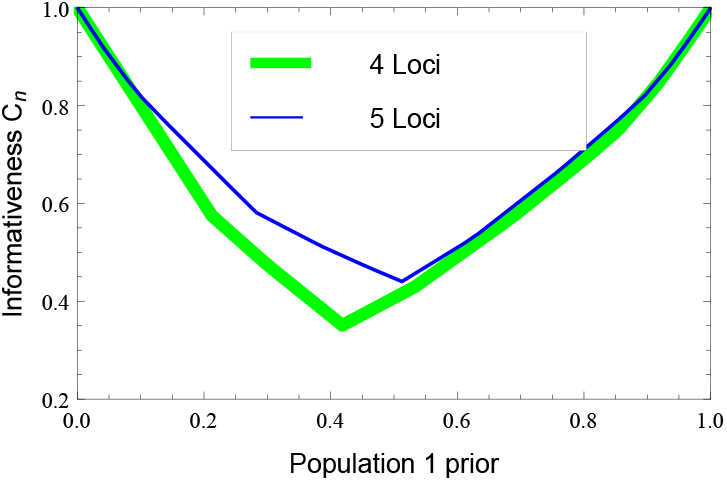
The prior-washout property: the effect of the prior on informativeness is diminished with more loci.

## Appendix D Proof the NC-based *C_n_* complies with a subset of the criteria

1. *Zero:* If *P* = *Q* (i.e., *p_i_* = *q_i_* for all *i*) then by definition *α* = 0.5 (since these are effectively the same population) and thus *h_k_* = *g_k_*, *D_k_* = 0 and *d_k_* = 0 for all *k*. From Eq. (9), and since the sum of all genotype frequencies from each single population is 1,

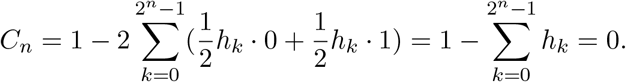 It remains to be proven in future work whether conversely, *C_n_* = 0 implies *P* = *Q* (i.e., *p_i_* = *q_i_* for all *i*) and *α* = 0.5.
2. *Performance:* This criterion fails. The discontinuity inherent in the classifier expression *d_k_* renders *C_n_* decreasing as *n* → *n* + 1 under certain scenarios (see Fig. D.1A).
3. *Asymptotics:* A formal proof that *C_n_* → 1 left for future work, but numerical simulations strongly indicate it (see Fig. D.1B for illustration).
4. *Neutrality:* We need to prove that if *p*_*n*+1_ = *q*_*n*+1_ then *C*_*n*+1_ = *C_n_*.

#### Proof.

From Eq. (4.2)we have,

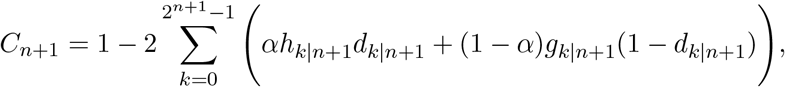

where the subscript *k*|*n* + 1 designates that the term is defined with reference to *n* +1 loci. First notice that the classifier expression does not change,

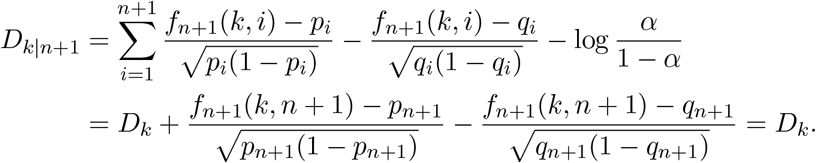

since *p*_*n*+1_ = *q*_*n*+1_ from assumption. Finally, we split the multiple sum into two multiple sums, the first in which genotypes have a ‘0’ allele as the *n* + 1 locus, and the second in which they have a ‘1’ allele at that locus, noticing also that the sum from 2^*n*^ to 2^*n*+1^ – 1 equals the sum from 0 to 2^*n*^ – 1 when the *n* + 1 locus is fixed,

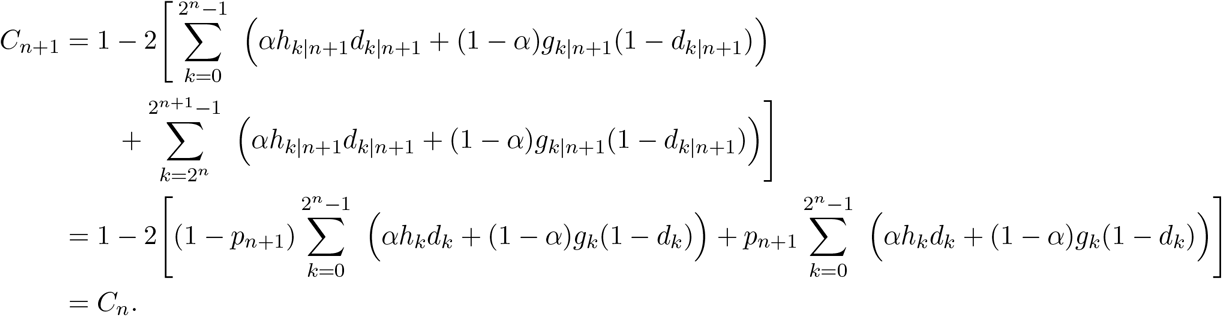
5. *Continuity:* This criterion fails (e.g., see in Fig. D.2 the discontinuity points).
6. *Dominance:* We need to prove that for any finite number of loci, *C_n_* → 1 iff |*q_i_ – p_i_*| → 1.

#### Proof.

We prove, without loss of generality, that if *q*_*n*+1_ → 1 and *p*_*n*+1_ → 0 then *C*_*n*+1_ → 1. In fact, we have

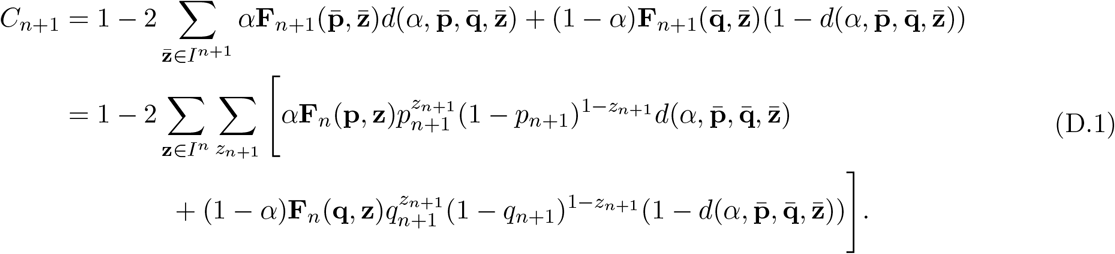

Note that the classifier 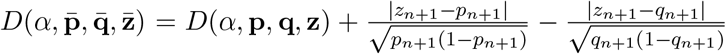. It implies that if *p*_*n*+1_ → 0 and *q*_*n*+1_ → 1 then

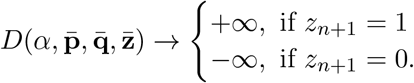

Therefore

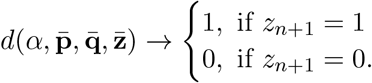

Thus, when *p*_*n*+1_ → 0 and *q*_*n*+1_ → 1 we have 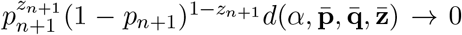 and 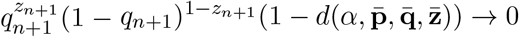. It follows from Eq. (D.1) that *C*_*n*+1_ → 1.
7. *Delta:* This criterion fails (e.g. see in Fig. D.2 the decrease of *C_n_* at several locations).
8. *Invariances:* [a] since *D_k_* is commutative as a sum, and *h_k_* and *g_k_* are commutative as a product, *C_n_* is invariant to different ordering of the loci; [b] *C_n_*(*α, P, Q*) = *C_n_*(1 – *α,Q,P*) follows from the symmetry of *D_k_* and *C_n_* with respect to *p_i_* and *q_i_*; [c] the simultaneous substitution of *p_i_* with (1 – *p_i_*) and *q_i_* with (1 – *q_i_*) simply changes the order of summation in *D_k_* and *C_n_*. We illustrate this with reference to *C*_1_, and this is easily proven by induction on *n*. We write down the explicit form of *C*_1_,

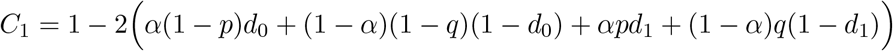

where,

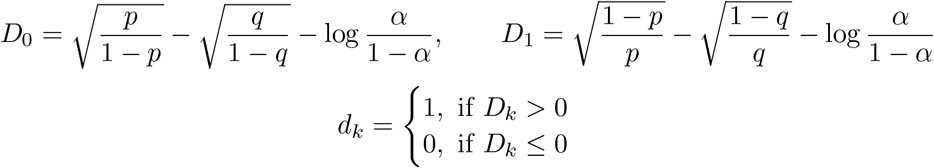 Denote by 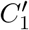 the measure following this substitution. It is straightforward that,

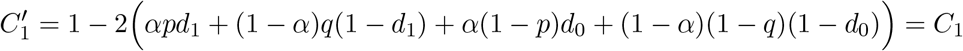

since *D_k_* (and *d_k_*) were similarly modified by the substitution,

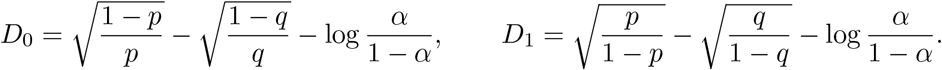
9. *Priors:* At the limit *α* → 0, *D_k_* → +∞, *d_k_* = 1 and thus in (9) both *αh_k_d_k_* → 0 and (1 – *α*)*g_k_*(1 – *d_k_*) → 0, therefore *C_n_* → 1. Similarly, at the limit *α* → 1, *D_k_* → −∞, = 0 and both *αh_k_d_k_* → 0 and (1 – *α*)*g_k_*(1 – *d_k_*) → 0, therefore *C_n_* A 1.

**Fig D.1:**
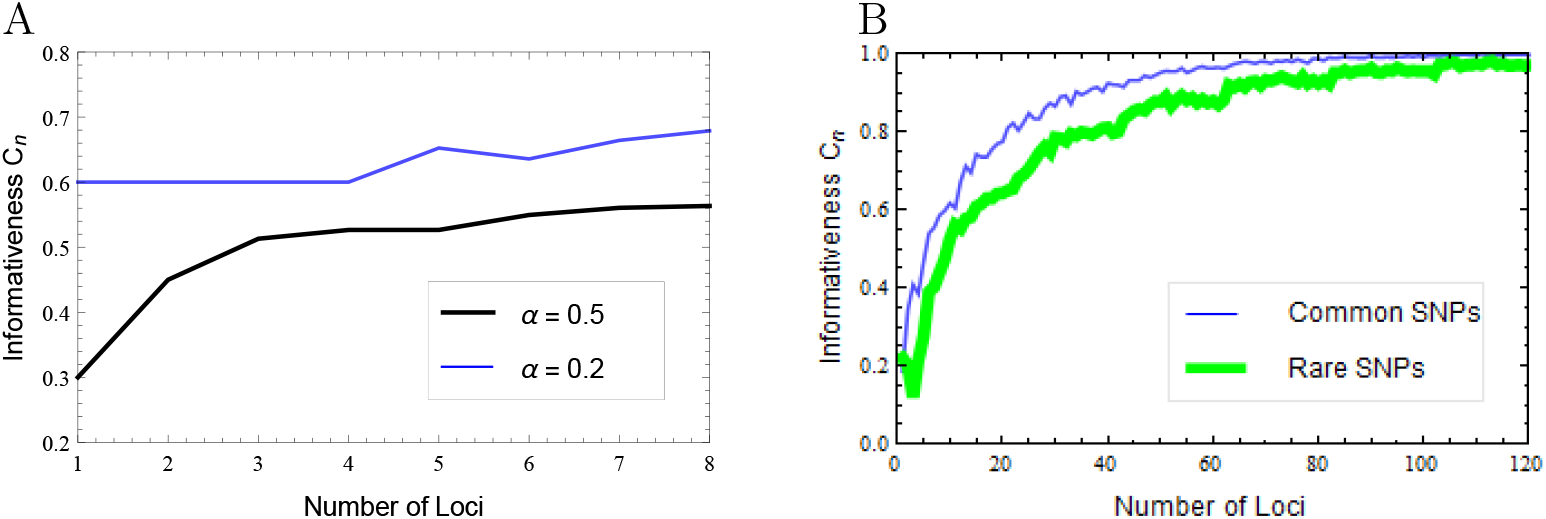
*C_n_* based on a NC classifier fails *performance* (here *p_i_* = 0.1/*q_i_* = 0.4 for the first four loci and subsequently *p_i_* = 0.5/*q_i_* = 0.62). | B: *C_n_* based on a NC classifier satisfies asymptotics criterion, here simulated with both common (*p_i_* = 0.2/*q_i_* = 0.45) and rare alleles (*p_i_* = 0.02/*q_i_* = 0.12) and *α* = 0.4.

**Fig D.2:**
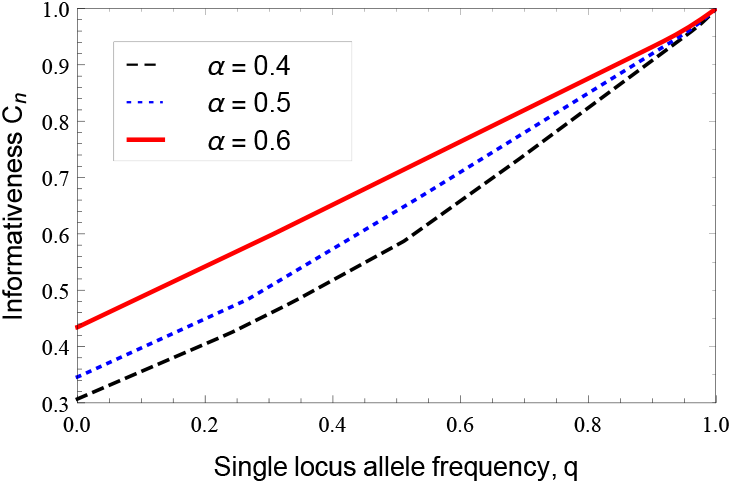
*C_n_* based on a NC classifier satisfies the *dominance* criterion but fails the *continuity* and *delta* criteria. Shown here are three cases corresponding to different class priors, for 5 loci where *q*_5_ changes from 0+ through 1– while *p*_5_ = 0+ the other 4 loci we given arbitrary frequencies: *p*_1_ = 0.01/*q*_1_ = 0.3,*p*_2_ = 0.2/*q*_2_ = 0.35, *p*_3_ = 0.4/*q*_3_ = 0.3, *p*_4_ = 0.1/*q*_4_ = 0.15.

## Appendix E Proof of compliance of Sampling-based *C_n,m_* with the new criteria

Here we prove the two new criteria and also show by simulation compliance with two other criteria from *C_n_*. We first rewrite Eq. (5.2) in the form of

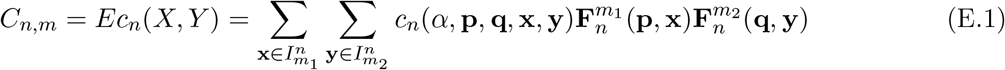

where

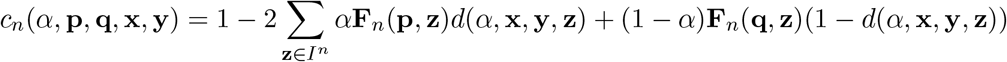

with

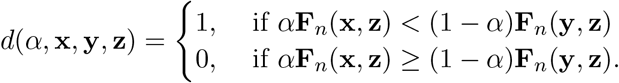

Note that we have immediately that *c_n_*(*α*, **p, q, p, q**) = *C_n_*.

[3] *Asymptotics:* A proof is beyond the scope of this paper, but numerical simulations validate it (see Fig. E.1A for illustration).

[6] *Dominance:* A formal proof is beyond the scope of this paper, but numerical simulations validate it (see Fig. E.1B for illustration).

[1*] *Sampling-Effect:* Need to show that for any sample size *m* > 0, *C_n,m_* ≤ *C_n_*. In fact, for given 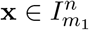, 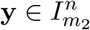, for each **z** ∈ *I^n^* we have *d*(*α*, **x, y, z**) ∈ {0,1}, therefore

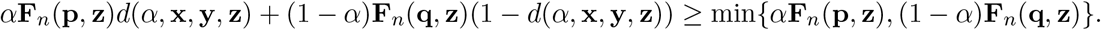

**Fig E.1:**
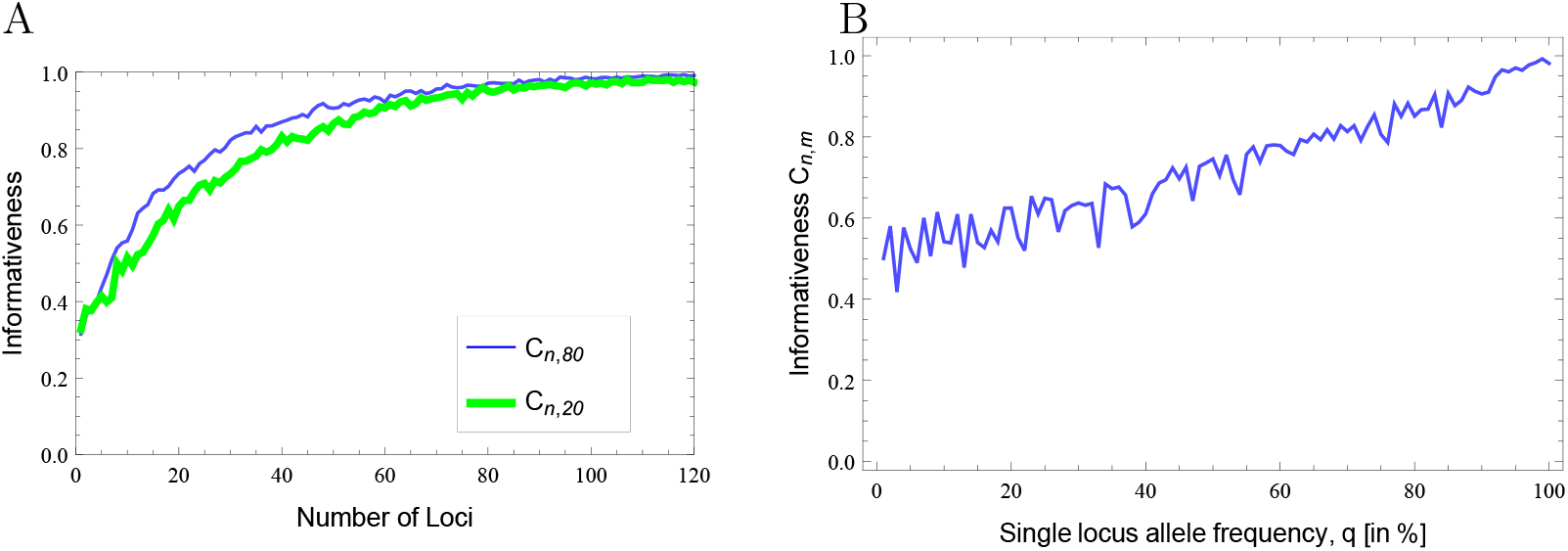
|**A**: *C_n,m_* satisfies the *asymptotics* criterion, simulated here for both 80 and 20 samples from each population, *α* = 0.4.|*B :C_n,m_* satisfies the *Dominance* criterion: *C_n,m_* → 1 as |*q_i_* – *p_i_*| → 1, simulated here for 20 samples, 8 loci, *α* = 0.7. In both examples we used *p_i_* = 0.1/*q_i_* = 0.3 (using Monte Carlo simulations).

It implies that, for given 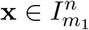, 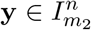, we have

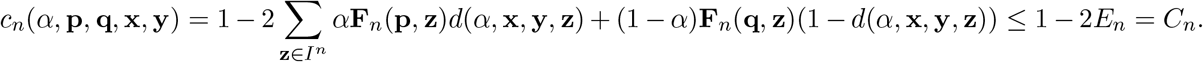

Thus

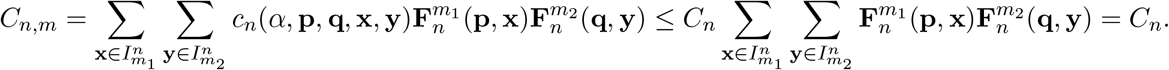

It completes the proof (see Fig.E.2A for illustration).

[2*] *Sampling-Convergence:* We need to show that as sample size increases *C_n,m_* → *C_n_*. In fact, we note that when *m*_1_ → ∞, 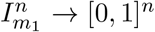 and 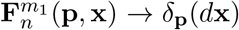. Similarly, when *m*_2_ → ∞, 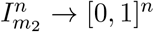 and 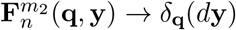. Therefore when *m* = {*m*_1_,*m*_2_} → ∞ we have

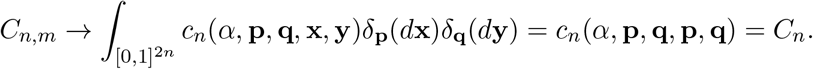

It completes the proof (see Fig. E.2B for illustration).

**Fig E.2:**
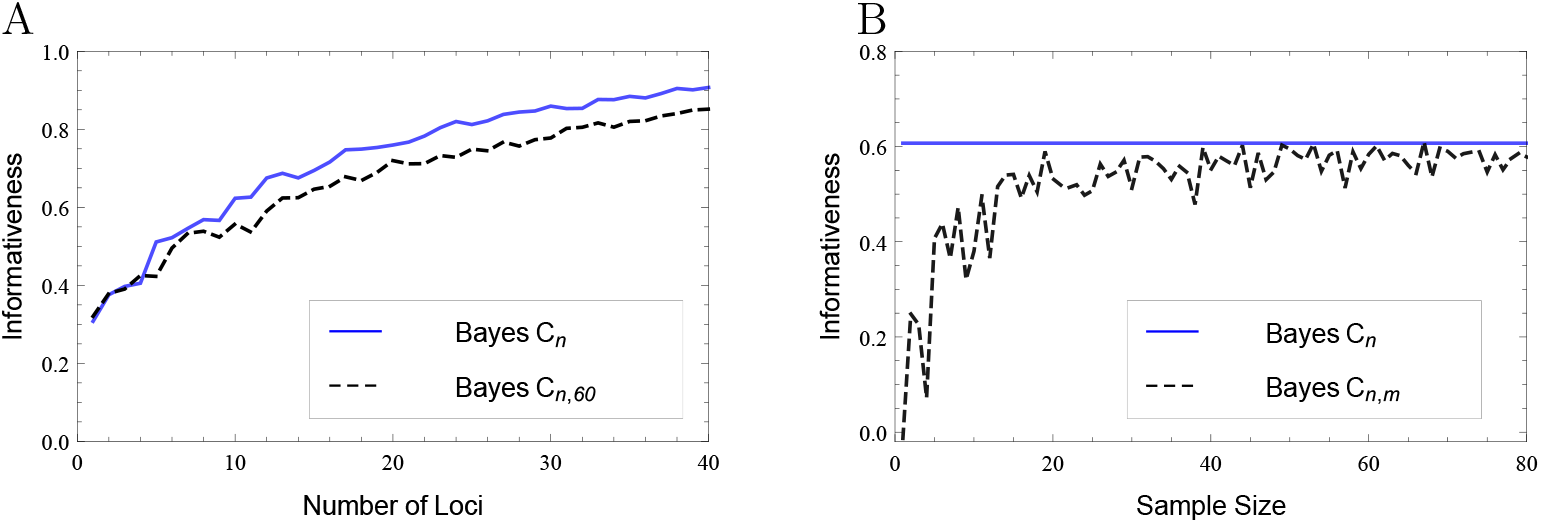
| A: *C_n,m_* satisfies the sampling-effect criterion. Shown here for 60 samples from each population, *α* = 0.6. | B: *C_n,m_* satisfies the sampling-convergence criterion. Shown here for 10 loci, where *α* = 0.4. In both examples we used *p_i_* = 0.1/*q_i_* = 0.3 (with Monte Carlo simulations).

## Appendix F Proof that every monotonic function of a compliant informativeness measure is also compliant

Given a function *f* : [0,1] → [0,1] such that

a. *f*(*x*) = 0 iff *x* = 0;
b. *f*(*x*) = 1 iff *x* = 1;
c. *f* is continuous;
d. *f* is monotone.

We show here that if *C_n_* is a compliant informativeness measure, then so is *f*(*C_n_*). In fact, the *Zero* criteria is followed from (a); the *Performance* criteria is followed from (d); the *Asymptotic* criteria is followed from (b) and (c); the *Neutrality* criteria is followed from the definition of a single-value function *f*; the *Continuity* criteria is followed from (c); the *Dominance* criteria is followed from (b) and (c); the *Delta* criteria is followed from (d); the *Invariances* criteria is followed from the definition of a single-value function *f*; the *Prior* criteria is followed from (b) and (c).

## Appendix G The alternative non-decision-theoretic formulation of 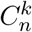 in compliance with all criteria

We construct here a family of measures 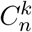, parametrized by the integer *k* : 1, … , ∞,

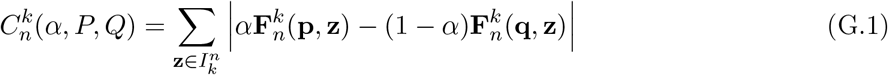

where **p** = (*p*_1_, … ,*p_n_*), **q** = (*q*_1_, … ,*q_n_*), 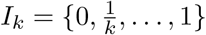 and

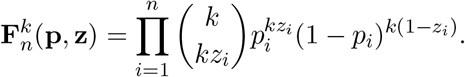

Note that 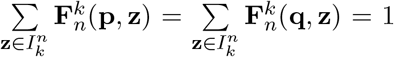, by using the same technique as in Appendix A, we also have

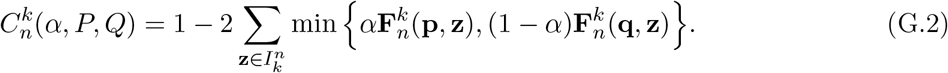

We show that these measures are compliant with all criteria.

1. *Zero:* (⇐): If *P* = *Q* (i.e. *p_i_* = *q_i_*, ∀*i* = 1, … , *n*) and *α* = 0.5 then we have 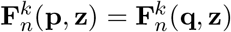 for all 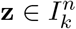. Thus from Eq. (G.1), and since the sum of all genotype frequencies from a single population is 1,

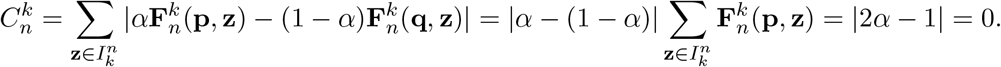

(⇐): Conversely, we need to show that 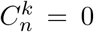 implies *P* = *Q* (i.e., *p_i_* = *q_i_*, ∀*i* = 1, … ,*n*) and *α* = 0.5. First, notice that if 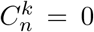 then trivially each summand of Eq. (G.1) must be zero, i.e. 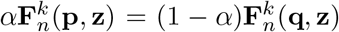, 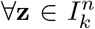. By summing for all 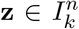 we have *α* = 1 – *α* or *α* = 0.5. Therefore 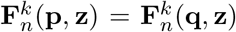, 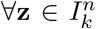. Denote by 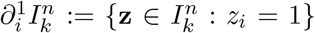. Then we have for every *i* = 1, … , *n*

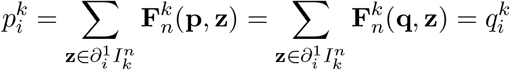

which implies *p_i_* = *q_i_*.
2. *Performance:* We need to prove that 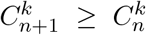. In fact, by denoting 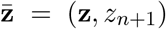, 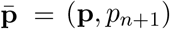, 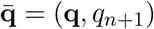, we have

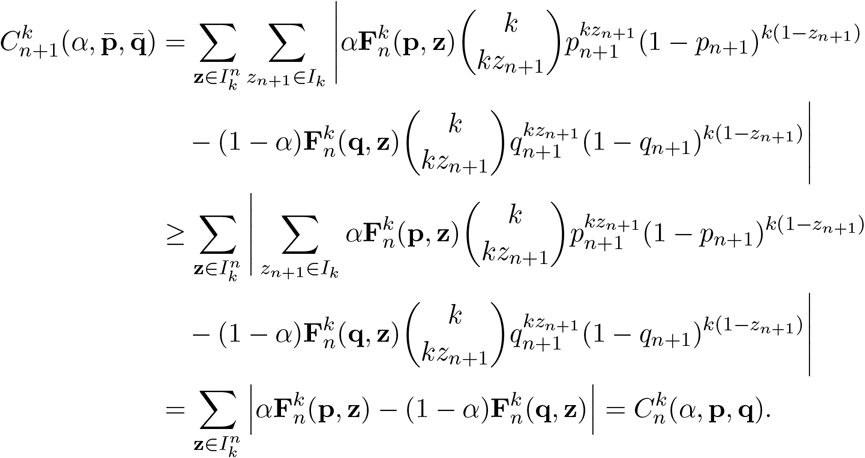
3. *Asymptotics:* First of all, note that

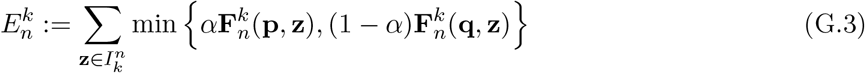

is also a Bayes error for classifying over two *n*-loci (*k* + 1)-allele populations. By applying the same technique as in [Tal, 2012b, Appendix B.1], 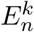 is bounded above by 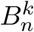, the misclassification rate from simple multinomial model. Moreover, we have,

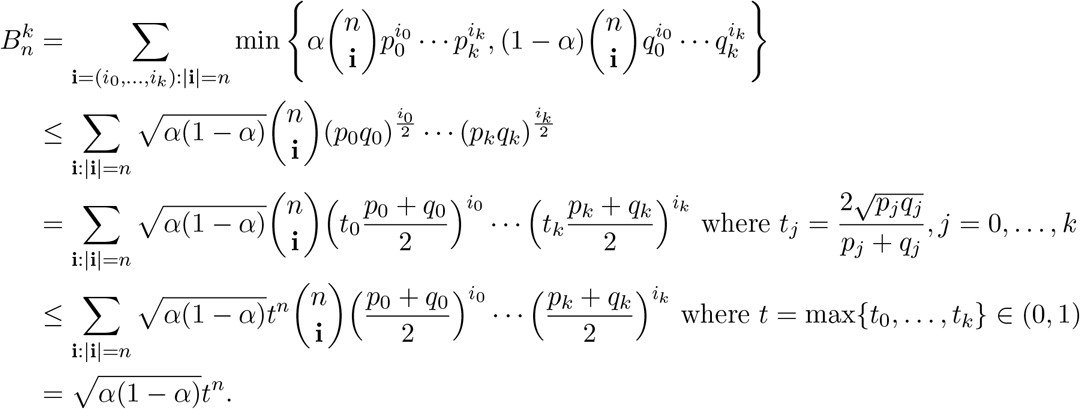 Thus, 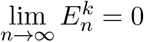. It implies 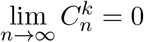.
4. *Neutrality:* We need to prove that 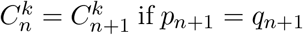 if *p*_*n*+1_ = *q*_*n*+1_. In fact, by denoting 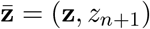, 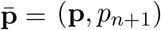, 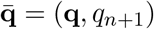, we have

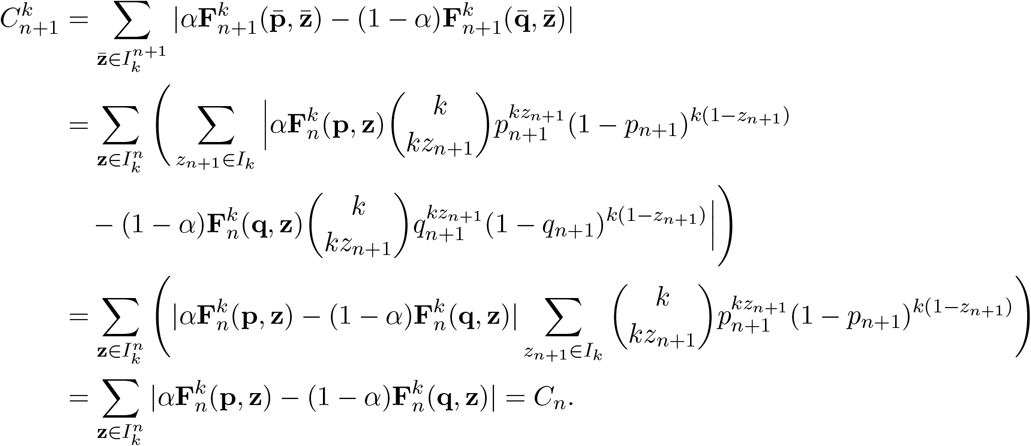
5. *Continuity:* From Eq. (G.1), 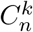 is a sum of absolute values of continuous functions and therefore has no singularities, and since *p_i_,q_i_* and *α* are real-valued parameters, 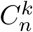 is continuous with respect to its parameters.
6. *Dominance:* (⇒) We need to prove that for a fixed n and some fixed *α* ∈ (0,1), if there exists *i* ∈ {1, … , *n*} such that |*q_i_* – *p_i_*| → 1 then 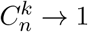. In fact, without loss of generality we assume that *p_n_* → 0 and *q_n_* → 1. We note that

- If *z_n_* = 0 then 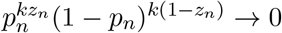 and then 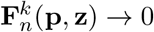
- If *z_n_* = 0 then 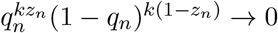 and then 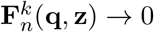. Therefore 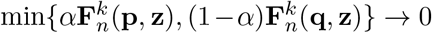 for all 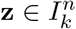. This results in 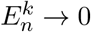 and consequently from Eq. (G.2) 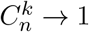. (⇐) Conversely, assume that 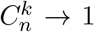 for a fixed *n* and some fixed *α* ∈ (0,1), we need to prove that there exists *i* ∈ {1, … , *n*} such that |*q_i_* – *p_i_*| → 1. In fact, we first note that **p** ≠ **q** because otherwise 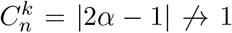 which is a contradiction. Moreover, because 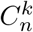 satisfy the neutrality and invariant criteria, we can assume that *p_i_* < *q_i_* for all *i*. Now, put 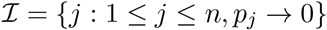 and 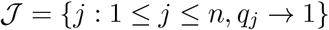. Because 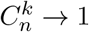, it implies

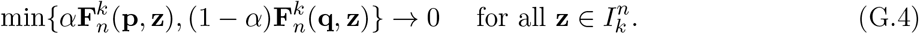 By choosing **z** = (0, … , 0) we imply 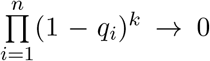, therefore 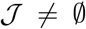. Similarly, by choosing **z** = (1, … , 1) we imply 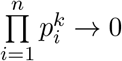, therefore 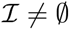. Assume that 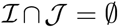 then we can choose 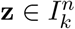 such that 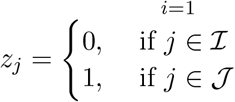. Note that

- 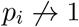 for 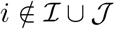 (otherwise, it implies *q_i_*(> *p_i_*) → 1 and contradicts to the definition of 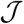);
- 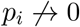 for 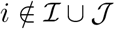 and 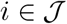 (from the definition of 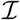). Therefore we have 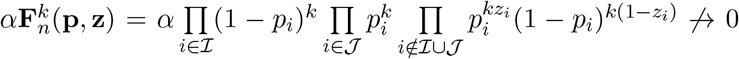. Similarly, we have 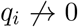 for 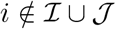; 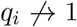 for 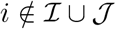 and 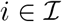 and therefore 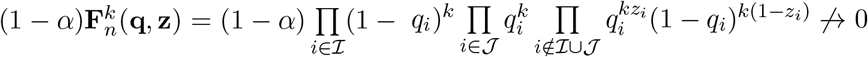. It implies that 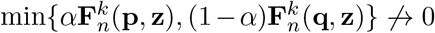 which contradicts to (G.4). Thus 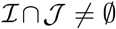 which implies the proof.
7. *Delta:*
8. *Invariances:* [a] since the genotype probabilities 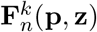 and 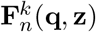 in Eq. (G.1) are each a commutative product of allele frequencies from all loci, 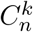 is invariant to different ordering of loci; [b] 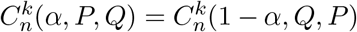 follows from the presence of an absolute value in the formulation of Eq. (G.1); [c] the simultaneous substitution of *p_i_* with (1 – *p_i_*) and *q_i_* with (1 – *q_i_*) simply changes the order of the summation terms in Eq. (G.1) and thus does not affect 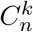.
9. *Prior:* If *α* → 0 or *α* → 1 then one of the two terms within the sum in Eq. (G.1) diminishes to zero and what remains in the limit is the sum over all genotype probabilities in one population, which equals 1, therefore 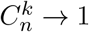.

## Appendix H Proof that the alternative 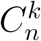 is not a monotonic function of the Bayes *C_n_*

More precisely, we show that, for given *n* and *k* > 1 we can find (*α*, **p, q**) and (*α*′, **p′, q′**) such that *C_n_*(*α*, **p, q**) = *C_n_*(*α*′, **p**′, **q**′) but 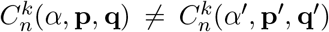. In fact, we can choose, for example, *α* = *α*′ = 0.5, *p*_1_ = 0.1, *q*_1_ = 0, 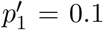, 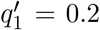, and *p_i_* = *q_i_* = 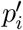 = 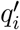 = 0.3 for all *i* = 2, … , *n*. By applying the *Neutrality* criteria, we have 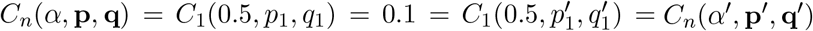 but 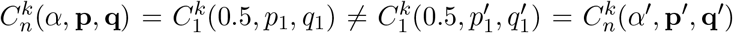. The figure Fig. H.1 illustrate the behavior of the difference 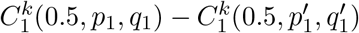 when *k* runs from 1 to 10.

**Fig H.1:**
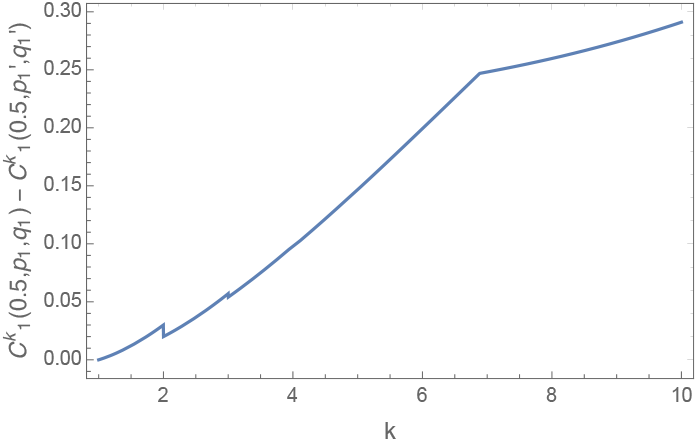
Behavior of 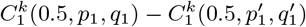

## Footnotes

1 A simple counter example for a loci pair is: *p*_1_ = 0.24/*q*_1_ = 0.18,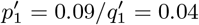 and prior *α* = 0.51.

2 e.g., *n* = 1, *p* = 1/4, *q* = 1/2, *r* = 3/4, *α* = 2/5, *β* = 3/5,7 = 1/2. In general, for higher dimensions such failure may occur with *q_i_* = (*p_i_r_i_*)^1/2^, *β* = 1 − *α, γ* =1/2; e.g., *n* = 3; *p_i_* = 0.7; *π* = 0.95; *q_i_* = (*p_i_r_i_*)^1/2^, *α* = 0.45, *β* = 0.55,7 = 1/2.

3 e.g., the 3rd locus here is uninformative: *p*_1_ = 0.02/*q*_1_ = 0.24, *p*_2_ = 0.2/*q*_2_ = 0.1,*p*_3_ = 0.1/*q*_3_ = 0.25 with *α* = 0.7.

4 It is also possible to use a proper Bayesian approach for estimating allele frequencies, with Beta priors at each SNP locus, as in [Rannala and Mountain, 1997, Eq. 5].

5 In simulating *C_n,m_* we replace allele frequency estimates of *zero* with a small constant, 1/(*m*+1), a common procedure to avoid zero genotype frequencies ([Rosenberg, 2005]; [Phillips et al., 2007]).

6 Note that the existence of this non-degenerate range around zero for *ε* does not depend on the allele frequency at the extra locus (denoted in this proof as *p*).

7 It is true almost surely since the probability that a genotype would have the same frequency in both populations, weighed by the priors, is vanishingly small (even with only one locus, since allele frequencies differ between populations).

## References

[Battiti, 1994] Battiti, R. (1994). Using mutual information for selecting features in supervised neural net learning. IEEE Transactions on Neural Networks, 5(4):537–550.

[Brown et al., 2012] Brown, G., Pocock, A., Zhao, M.-J., and Luján, M. (2012). Conditional likelihood maximisation: A unifying framework for information theoretic feature selection. J. Mach. Learn. Res., 13(1):27–66.

[Carja and Feldman, 2012] Carja, O. and Feldman, M. W. (2012). An equilibrium for phenotypic variance in fluctuating environments owing to epigenetics. Journal of The Royal Society Interface, 9(69):613–623.

[Cornuet et al., 1999] Cornuet, J.-M., Piry, S., Luikart, G., Estoup, A., and Solignac, M. (1999). New methods employing multilocus genotypes to select or exclude populations as origins of individuals. Genetics, 153(4):1989–2000.

[Cover and Thomas, 2006] Cover, T. M. and Thomas, J. A. (2006). Elements of Information Theory. Wiley.

[Csiszár, 2008] Csiszár, I. (2008). Axiomatic characterizations of information measures. Entropy, 10(3):261–273.

[Degen et al., 2017] Degen, B., Blanc-Jolivet, C., Stierand, K., and Gillet, E. (2017). A nearest neigh-bour approach by genetic distance to the assignment of individual trees to geographic origin. Forensic Science International: Genetics, 27:132–141.

[Delaigle and Hall, 2012] Delaigle, A. and Hall, P. (2012). Achieving near perfect classification for functional data. Journal of the Royal Statistical Society: Series B (Statistical Methodology), 74(2):267–286.

[Ding et al., 2011] Ding, L., Wiener, H., Abebe, T., Altaye, M., Go, R. C., Kercsmar, C., Grabowski, G., Martin, L. J., Khurana Hershey, G. K., Chakorborty, R., and Baye, T. M. (2011). Comparison of measures of marker informativeness for ancestry and admixture mapping. BMC Genomics, 12:622–622.

[Dudoit et al., 2002] Dudoit, S., Fridlyand, J., and T.P., S. (2002). Comparison of discrimination methods for the classification of tumors using gene expression data. Journal of the American Statistical Association, 97(457):77–87.

[Edwards, 2003] Edwards, A. (2003). Human genetic diversity: Lewontin’s fallacy. BioEssays, 25(8):798–801.

[Estoup and Angers, 1998] Estoup, A. and Angers, B. (1998). Microsatellites and minisatellites for molecular ecology : theoretical and empirical considerations. Advances in Molecular Ecology, pages 55–86. (ed. Carvalho G.), NATO press.

[Gattepaille and Jakobsson, 2012] Gattepaille, L. M. and Jakobsson, M. (2012). Combining markers into haplotypes can improve population structure inference. Genetics, 190(1):159–174.

[Grall-Maes and Beauseroy, 2002] Grall-Maes, E. and Beauseroy, P. (2002). Mutual information-based feature extraction on the time-frequency plane. IEEE Transactions on Signal Processing, 50(4):779–790.

[Hastie et al., 2009] Hastie, T., Tibshirani, R., and Friedman, J. (2009). The elements of statistical learning. Springer Series in Statistics. Springer, New York, second edition. Data mining, inference, and prediction.

[Huang and Chow, 2005] Huang, D. and Chow, T. W. S. (2005). Effective feature selection scheme using mutual information. Neurocomput., 63:325–343.

[Huang and Rong, 2009] Huang, J. and Rong, P. (2009). A Hybrid Genetic Algorithm for Feature Selection Based on Mutual Information, pages 125–152. Springer US, Boston, MA.

[Khosravifard et al., 2007] Khosravifard, M., Fooladivanda, D., and Gulliver, T. (2007). Confliction of the convexity and metric properties in f-divergences. IEICE Trans. on Fundamentals, 9:1848–1853.

[Last et al., 2001] Last, M., Kandel, A., and Maimon, O. (2001). Information-theoretic algorithm for feature selection. Pattern Recogn. Lett., 22(6-7):799–811.

[Lawson et al., 2012] Lawson, D. J., Hellenthal, G., Myers, S., and Falush, D. (2012). Inference of population structure using dense haplotype data. PLOS Genetics, 8(1):1–16.

[Liao et al., 2009] Liao, H., Liu, Y., and Michael, K. (2009). Shrunken dissimilarity measure for genome-wide snp data classification. Technical report, The Third International Symposium on Optimization and Systems Biology (OSB’09). pp. 73–80.

[Meila, 2007] Meila, M. (2007). Comparing clusterings-an information based distance. Journal of Multivariate Analysis, 98(5):873–895.

[Nguyen et al., 2009] Nguyen, X., Wainwright, M. J., and Jordan, M. I. (2009). On surrogate loss functions and f-divergences. Ann. Statist., 37(2):876–904.

[Patterson et al., 2006] Patterson, N., Price, A., and Reich, D. (2006). Population structure and eigenanalysis. PLoS Genetic, 2(12):e190.

[Peng et al., 2005] Peng, H., Long, F., and Ding, C. (2005). Feature selection based on mutual information criteria of max-dependency, max-relevance, and min-redundancy. IEEE Transactions on Pattern Analysis and Machine Intelligence, 27(8):1226–1238.

[Phillips et al., 2007] Phillips, C., Salas, A., Sanchez, J., Fondevila, M., Gomez-Tato, A., Alvarez-Dios, J., Calaza, M., Casares de Cal, M., Ballard, D., Lareu, M., and Carracedo, A. (2007). Inferring ancestral origin using a single multiplex assay of ancestry-informative marker SNPs. Forensic Science International: Genetics, 1(3-4):273–280.

[Pritchard et al., 2000] Pritchard, J. K., Stephens, M., and Donnelly, P. (2000). Inference of population structure using multilocus genotype data. Genetics, 155(2):945–959.

[Rannala and Mountain, 1997] Rannala, B. and Mountain, J. L. (1997). Detecting immigration by using multilocus genotypes. Proceedings of the National Academy of Sciences, 94(17):9197–9201.

[Rodin, 2014] Rodin, A. (2014). Axiomatic method and category theory, volume 364 of Synthese Library. Studies in Epistemology, Logic, Methodology, and Philosophy of Science. Springer, Cham.

[Rosenberg, 2005] Rosenberg, N. A. (2005). Algorithms for selecting informative marker panels for population assignment. Journal of Computational Biology, 12(9):1183–1201.

[Rosenberg et al., 2003] Rosenberg, N. A., Li, L. M., Ward, R., and Pritchard, J. K. (2003). Informativeness of genetic markers for inference of ancestry. American Journal of Human Genetics, 73(6):1402–1422.

[Sampson et al., 2011] Sampson, J. N., Kidd, K. K., Kidd, J. R., and Zhao, H. (2011). Selecting snps to identify ancestry. Annals of Human Genetics, 75(4):539–553.

[Shannon, 1948] Shannon, C. E. (1948). A mathematical theory of communication. The Bell System Technical Journal, 27(3):379–423.

[Steuer et al., 2002] Steuer, R., Kurths, J., Daub, C. O., Weise, J., and Selbig, J. (2002). The mutual information: Detecting and evaluating dependencies between variables. Bioinformatics, 18(suppl2) : S231– –S240.

[Tal, 2012a] Tal, O. (2012a). The cumulative effect of genetic markers on classification performance: Insights from simple models. Journal of Theoretical Biology, 293:206–218.

[Tal, 2012b] Tal, O. (2012b). Towards an information-theoretic approach to population structure. In Turing-100. The Alan Turing Centenary, volume 10 of EasyChair Proceedings in Computing, pages 353–369. EasyChair.

[Tal, 2013] Tal, O. (2013). Two complementary perspectives on inter-individual genetic distance. Biosystems, 111(1):18–36.

[Tal et al., 2017] Tal, O., Tran, T. D., and Portegies, J. (2017). From typical sequences to typical genotypes. Journal of Theoretical Biology, 419:159–183.

[Tibshirani et al., 2002] Tibshirani, R., Hastie, T., Narasimhan, B., and Chu, G. (2002). Diagnosis of multiple cancer types by shrunken centroids of gene expression. Proceedings of the National Academy of Sciences, 99(10):6567–6572.

[Tibshirani et al., 2003] Tibshirani, R., Hastie, T., Narasimhan, B., and Chu, G. (2003). Class prediction by nearest shrunken centroids, with applications to dna microarrays. Statist. Sci., 18(1):104–117.

[Wang, 2006] Wang, J. (2006). Informativeness of genetic markers for pairwise relationship and relatedness inference. Theoretical Population Biology, 70(3):300–321.

[Witherspoon et al., 2007] Witherspoon, D. J., Wooding, S., Rogers, A. R., Marchani, E. E., Watkins, W. S., Batzer, M. A., and Jorde, L. B. (2007). Genetic similarities within and between human populations. Genetics, 176(1):351–359.

[Zhao et al., 2013] Zhao, M.-J., Edakunni, N., Pocock, A., and Brown, G. (2013). Beyond fano’s inequality: Bounds on the optimal f-score, ber, and cost-sensitive risk and their implications. J. Mach. Learn. Res., 14(1):1033–1090.

